# Single-Cell Electrophysiology Reveals Verapamil’s Disruption of Bacterial Membrane Energetics

**DOI:** 10.64898/2026.05.31.729094

**Authors:** Anaïs Biquet-Bisquert, Amélie Astezan, Malo Marmol, Peter L. Voyvodic, Neha Mohite, Francesco Pedaci, Ashley L. Nord

## Abstract

Verapamil, a clinically used calcium channel blocker, enhances the activity of several tuberculosis antibiotics, but its mechanism of action and physiological effects on bacteria remain unresolved. A central debate concerns whether verapamil primarily inhibits efflux pumps or disrupts membrane energetics. Here, we use *Escherichia coli* as a model system to quantify single-cell and population-level physiological responses to verapamil with high temporal resolution. Real-time measurements of the rotational speed of individual flagellar motors, a single-cell proxy for the proton motive force (PMF), reveal a heterogeneous response to verapamil: treated cells exhibit either a dose-dependent gradual decrease in PMF, or a rapid collapse of PMF. Although loss of the outer-membrane efflux channel TolC increases growth inhibition by verapamil, it does not alter the rapid PMF disruptions observed at the single-cell level, suggesting that efflux contributes to long-term susceptibility but not to the initial PMF disruption. Independent assays of population-level motility, pH, and membrane-integrity suggest that verapamil may selectively dissipate the electrical component of PMF while leaving intracellular pH largely unchanged. A minimal electrical circuit model captures both steady-state and dynamic behavior. Together, these findings demonstrate that verapamil rapidly and reversibly perturbs bacterial membrane energetics through a mechanism distinct from classical protonophores, helping to reconcile conflicting interpretations of its activity and clarifying how membrane effects may interact with efflux inhibition during antibiotic potentiation.

## 1 Introduction

Tuberculosis (TB), caused by the pathogenic bacterium *Mycobacterium tuberculosis* (Mtb), remains one of the deadliest infectious diseases in the world, causing an estimated 1.25 million deaths in 2023 [1]. Despite the availability of effective anti-TB drugs, treatment remains long and complex, typically requiring a multi-drug regimen administered over several months. This prolonged duration imposes a significant burden on patients and healthcare systems, complicates compliance, and contributes to the emergence of drug resistance. The growing prevalence of resistance to nearly every current anti-TB drug underscores the urgent need for therapeutic innovations, including strategies to improve the potency of existing antibiotics.

In recent years, the re-purposing of existing drugs to act synergistically with anti-TB agents has emerged as a promising approach [2, 3]. One such compound is verapamil, an FDA-approved calcium channel blocker widely used to treat cardiovascular disease [4]. Multiple studies have shown that verapamil potentiates the activity of several anti-TB drugs, both *in vitro* and *in vivo*, including rifampicin, bedaquiline, and clofazimine [5–11]. This potentiating effect appears to be independent of its Ca^2+^ channel antagonist activity [9]. The most widely accepted mechanism is that verapamil inhibits mycobacterial drug efflux pumps (Fig. 1), which are induced under stress conditions and in the macrophage environment [7, 9], where *M. tuberculosis* survives and reproduces within host macrophages, encountering nutrient limitation, acidic pH, and oxidative stress[7]. Efflux inhibition would increase the intracellular concentration of antibiotics, thereby restoring or amplifying their efficacy. Structural studies on related bacterial transporters support the plausibility of direct blockade of substrate-binding pockets by verapamil-like compounds [12]. Consistent with an efflux-based mechanism, verapamil has been shown to reduce rifampicin efflux in direct whole-cell assays in *M. tuberculosis* [11], and increased accumulation of intra-cellular rifampicin in its presence has also been reported [5]. Recent work demonstrated that verapamil and its metabolite norverapamil directly inhibit the MmpS5–MmpL5 efflux pump in Mycobacterium tuberculosis, thus potentiating bedaquiline activity both *in vitro* and during macrophage infection [13]. Additionally, verapamil has been proposed to act indirectly by increasing plasma drug levels by inhibiting mammalian transporters, such as P-glycoprotein, thus improving drug bioavailability [10, 14].

**Figure 1.**
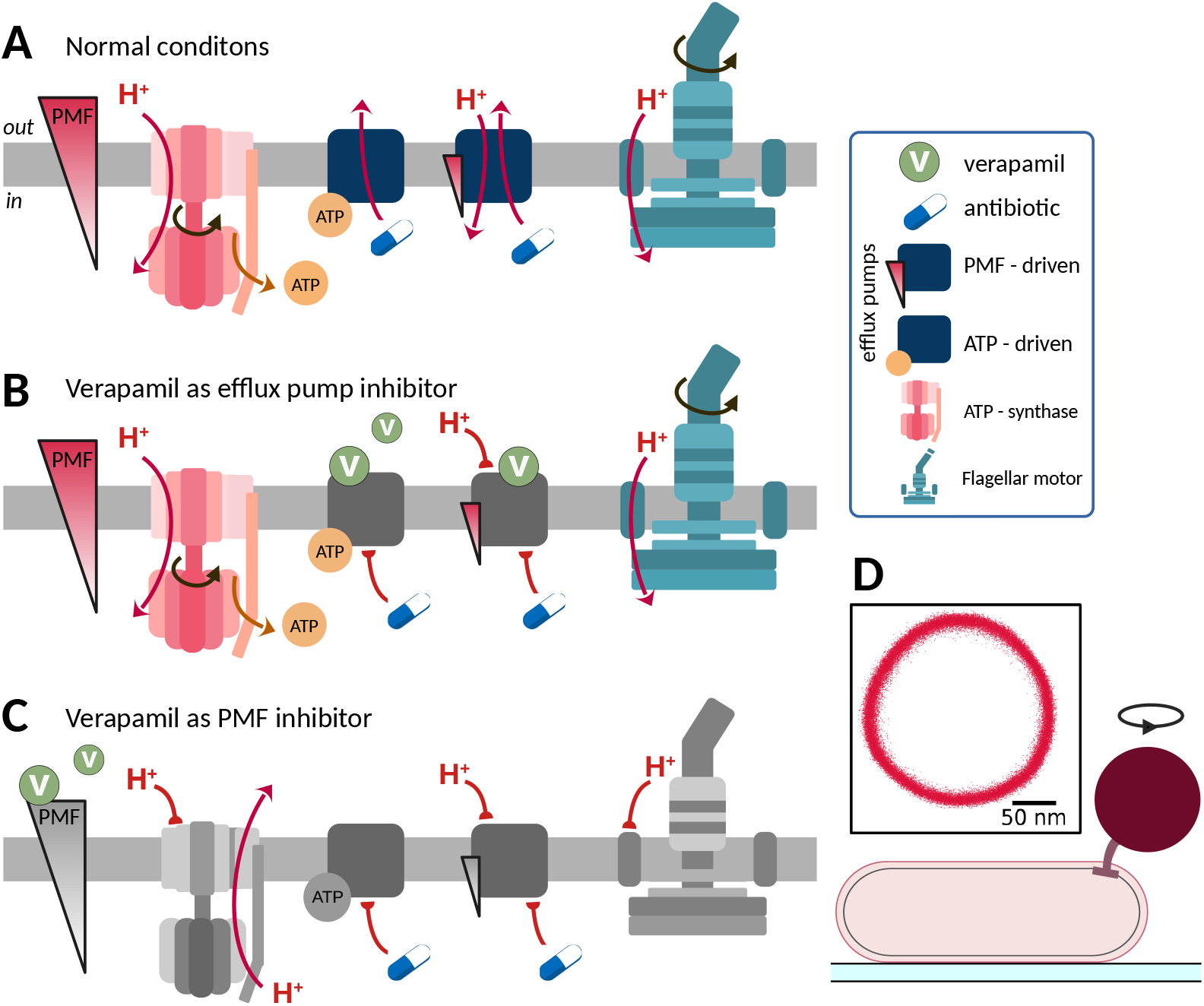
Competing models for how verapamil increases intracellular antibiotic accumulation, and a single-cell assay used to examine their impact on membrane energetics. (A) Under normal conditions, the proton motive force (PMF) drives ATP synthase, the bacterial flagellar motor, and efflux pumps, either directly or indirectly. Secondary efflux pumps (typically proton- or sodium-driven) harness the PMF directly, whereas primary (ATP-driven) pumps are powered indirectly via ATP generated by PMF-dependent ATP synthase activity. (B) Verapamil inhibits efflux pumps, either primary (ATP-driven) or secondary (ion-driven), by physically binding to the transporters and blocking the energy-dependent conformational changes required for drug export. (C) Verapamil disrupts the PMF, reducing ATP production and collapsing ion gradients, thereby impairing both ATP-driven and PMF-driven efflux pumps, as well as the bacterial flagellar motor, all of which depend directly or indirectly on membrane energetics. (D) Schematic of the bead assay. An *E. coli* cell was immobilized on a coverslip, and a bead was attached to its truncated flagellum. The inset shows a representative bead trajectory over several turns, from which rotation speed and thus motor output can be quantified.

However, several studies have challenged this efflux-centric view, instead proposing that verapamil may act by disrupting bacterial membrane energetics (Fig. 1C). Chen *et al*. found no evidence of increased intracellular rifampicin accumulation after verapamil treatment, either in isolated *M. tuberculosis* or infected macrophages [14]. Instead, they observed a rapid concentration-dependent collapse of both the membrane potential and the pH gradient, based on DiOC_2_(3) and acridine orange assays, respectively, suggesting that verapamil acts primarily by disrupting bacterial energetics. However, Ramakrishnan *et al*. recently revisited these measurements using TPP^+^ to probe membrane potential and acridine orange and benzoic acid uptake to assess the pH gradient[13]. At 100 μM verapamil, they observed partial dissipation of the pH gradient with acridine orange, but no effect with benzoic acid and no detectable change in membrane potential, leading them to conclude that verapamil does not measurably perturb bacterial energetics under physiologically relevant conditions. Additionally, Lake *et al*. found that verapamil reduced rifampicin efflux in a concentration-dependent manner [11], contradicting the conclusion that intracellular drug accumulation is not affected. They also questioned the use of DiSC_3_(5) to assess membrane potential, noting that verapamil may interfere with dye binding, and that CCCP, used as a positive control, did not behave as expected in Chen *et al*.’s experiments. These discrepancies may, in part, reflect the known limitations of voltage-sensitive dyes, including their slow equilibration kinetics, susceptibility to efflux, and the potential for the dyes themselves to perturb the membrane potential [15–19].

Nonetheless, the biophysical plausibility of energetic disruption is supported by verapamil’s amphiphilic, membrane-partitioning nature [20], its ability to alter bilayer structure and dipole potential [21], its activation of envelope stress responses in both *E. coli* and *M. tuberculosis* [14, 22], and by recent work showing that a structurally optimized phenylalkylamine analog disrupts mycobacterial membrane integrity and bioenergetics while augmenting bedaquiline activity [23]. Although verapamil has a high pKa (*∼*8.9) and is therefore predominantly protonated under physiological conditions, which would be expected to limit passive diffusion across lipid bilayers [11, 24], multiple studies have nonetheless shown that it partitions and accumulates in bacterial membranes, consistent with its high lipophilicity and favorable membrane-partitioning properties [14, 20–22, 25].

Given these technical limitations and conflicting interpretations, alternative methods are required to assess the physiological impact of verapamil on bacterial energetics. A powerful approach involves single-cell measurements of the rotational velocity of the bacterial flagellar motor (BFM), a molecular machine driven by the proton motive force (PMF) across the cytoplasmic membrane [26]. Because the BFM speed is linearly proportional to the PMF in the low viscous load regime [26, 27], it provides a high-resolution real-time readout of changes in membrane energetics [28–31]. In this work, we use BFM-based electrophysiology to quantify the dose-dependent effects of verapamil on PMF in single *E. coli* cells, explore heterogeneity in these responses, and investigate the potential involvement of efflux systems using a Δ*tolC* mutant. Our results reveal a dose-dependent reduction in PMF and pronounced heterogeneity in single-cell responses, providing new insight into how verapamil perturbs bacterial energetics and inform its use as an adjunct in TB therapy.

## 2. Results

### 2.1 Heterogeneity in single cell response to Verapamil

To quantify verapamil-induced changes in PMF, we performed high-resolution single-cell recordings of *E. coli* flagellar motor rotation (Fig. 1D; see Materi-als and Methods), using motor speed as a real-time readout of membrane energetics [27, 28, 30, 32–34].

The PMF is defined as:

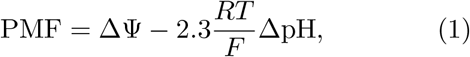

where ΔΨ *≡* Ψ_int_ − Ψ_ext_ is the transmembrane electric potential, and ΔpH *≡* pH_int_ − pH_ext_ is the pH gradient across the membrane. Here, the subscript *int* refers to the cytoplasmic (intracellular) side of the membrane, and *ext* refers to the periplasmic or extracellular side. *F* is the Faraday constant, *R* is the gas constant, and *T* is the absolute temperature (2.3*RT/F ≃* 60 mV at 300 K). Under aerobic conditions, wild-type *E. coli* typically maintains a PMF of approximately −150 mV, the negative sign reflects the inward directed electrochemical driving force on protons [35].

We tested the effects of verapamil across a large range of concentrations from 50 μM to 1 mM (Fig. 2A). In these experiments, verapamil solutions were flowed through a microfluidic chamber containing surface-adhered *E. coli*, and the rotation of individual flagellar motors was monitored before, during, and after verapamil exposure. Motor rotational speeds were measured using a bead assay, in which the rotation of a microbead attached to a sheared flagellum is tracked (see Materials and Methods and Fig. 1D). We observed a systematic, dose-dependent decrease in the average motor speed across the population (Fig. 2C), suggesting that verapamil decreases the PMF in a concentration-dependent manner.

**Figure 2.**
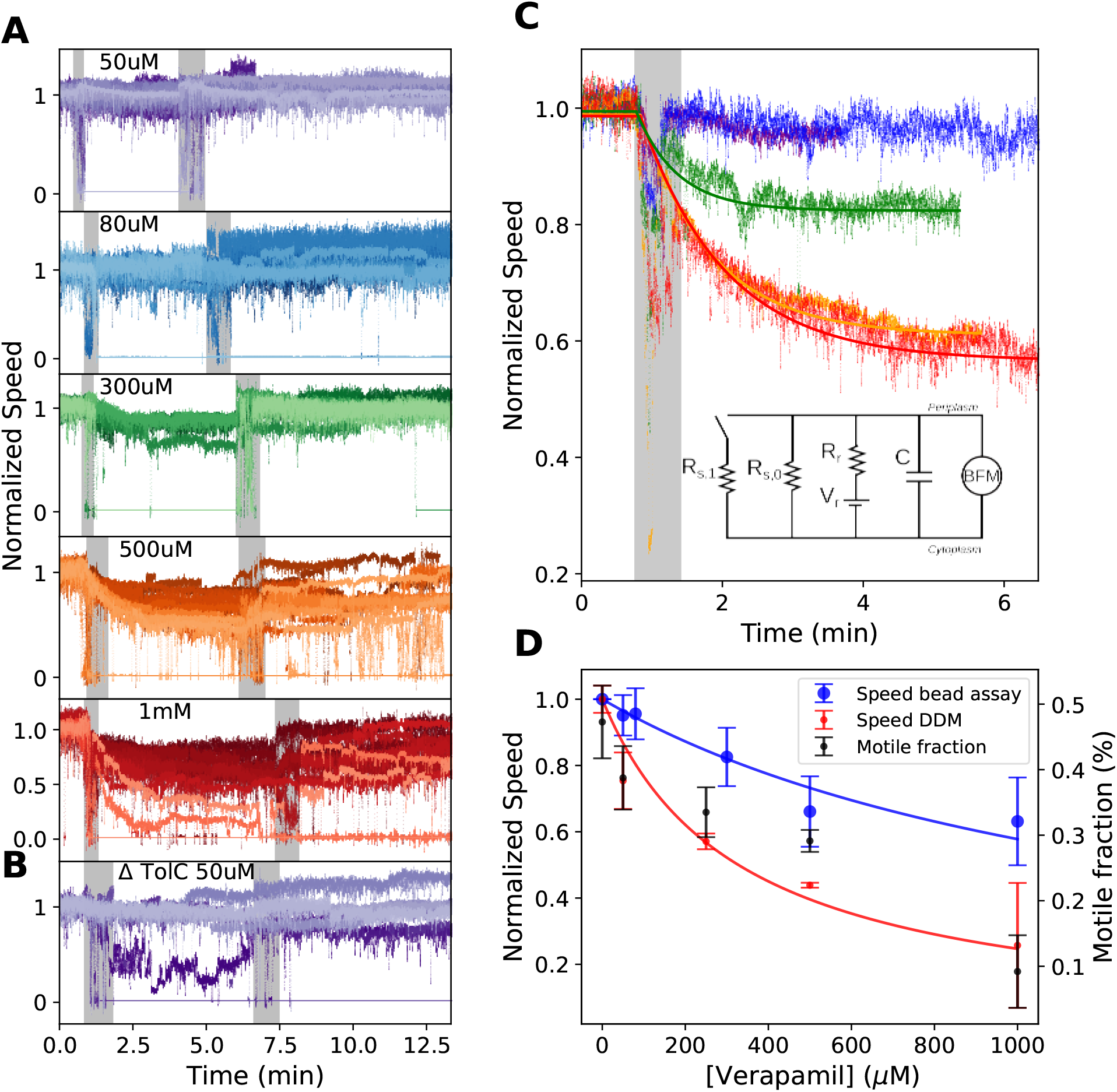
Verapamil induces a dose-dependent decrease in BFM speed. (A) Effect of verapamil on BFM speed at five concentrations: 50 μM, 80 μM, 300 μM, 500 μM, and 1 mM. Motors initially rotate in motility buffer (MB). Verapamil was introduced into the flow channel at approximately *t* = 50 s (first grey bar) and washed out at approximately *t* = 5 min (second grey bar). Speed was normalized by dividing by the mean speed during the initial period in MB prior to verapamil addition. Each trace corresponds to an individual motor (*n* = 22, 13, 9, 24, and 33 motors for increasing concentrations, respectively). (B) Effect of verapamil on Δ*tolC* motor speeds at 50 μM (*n* = 9 motors). (C) Average normalized speed of motors from (A) that continued rotating (excluding those that stopped). Thick lines indicate exponential fits, yielding characteristic time constants *τ*_300_ = 33.8 s, *τ*_500_ = 59.2 s, and *τ*_1000_ = 73.9 s. Inset: Equivalent circuit model. *R*_*s*,0_ represents membrane proton sink resistance prior to verapamil addition, modified by an additional pathway *R*_*v*_ . *V*_*r*_ and *R*_*r*_ denote the respiration-associated voltage source and internal resistance, respectively, *C* is the membrane capacitance, and the BFM is modeled as a voltmeter proportional to membrane potential. (D) Comparison of single-motor (bead assay) and population-level (DDM) measurements. Blue: steady-state BFM speed (bead assay) as a function of verapamil concentration, computed over a 30 s window after 4 min of exposure. The blue curve shows a hyperbolic fit to Eq. 5 (*R*^2^ = 0.92). Red: normalized swimming speed measured by DDM. The right axis (black) shows the fraction of motile cells *α*. For DDM measurements and *α, n* = 3 samples per condition. Error bars represent standard deviations across individual motors (blue) and samples (red, black).

In addition to this trend, we also observed significant cell-to-cell heterogeneity. From 50 μM, two distinct classes of motor behavior emerged: most motors progressively slowed as verapamil concentration increased, while a smaller subpopulation exhibited an abrupt loss of rotation, consistent with a collapse of the PMF (Fig. 2A). Both responses were often observed simultaneously within the same field of view (see Supplementary Fig. S1), suggesting a true biological heterogeneity between individual cells. Moreover, we observed no correlation between initial motor speed and motor collapse (see Supplementary Fig. S1). Upon washout of verapamil, motor speed typically recovered to baseline for concentrations up to 300 μM, while recovery was often incomplete at higher concentrations (see Supplementary Fig. S2).

### 2.2 Efflux contributes to verapamil resistance but not to single-cell heterogeneity

The heterogeneous motor responses that we observe in response to verapamil treatment, where some motors gradually slow and others abruptly stop, are reminiscent of the effects reported for other membrane-active compounds. Le *et al*. found that treatment of *E. coli* with protonophores such as CCCP and indole resulted in bimodal distributions of growth, substrate transport, and flagellar motor speed at intermediate concentrations [32]. In particular, single-cell flagellar motor measurements showed that while some cells exhibited a decreased rotational speed (indicating a partial loss of PMF), others showed a complete motor stop (indicating full PMF collapse), akin to the behavioral classes we observed with verapamil. Le *et al*. showed that this heterogeneity is mediated by active efflux. They observed that in a Δ*tolC* mutant which lacked the outer membrane channel required for AcrAB–TolC-mediated export, the CCCP-induced heterogeneity was abolished. Instead, all cells showed uniform, gradual decreases in motor speed with increasing CCCP concentrations. This suggested that efflux activity contributes to a positive feedback loop wherein cells with strong initial PMF can power efflux pumps to expel protonophores, thereby preserving their PMF and maintaining efflux function. In contrast, cells with weaker PMF lose efflux capacity more rapidly, accelerating PMF collapse. TolC dependent drug efflux systems extrude not only CCCP, but also a broad spectrum of structurally diverse compounds, including antibiotics, dyes, and amphiphilic molecules, [36, 37], and deletion of *tolC* increases bacterial susceptibility to multiple antibiotics [38].

To test whether TolC similarly influences heterogeneity in the verapamil response, we first compared growth of wild-type and Δ*tolC* strains across a range of verapamil concentrations. The Δ*tolC* mutant was significantly more sensitive: while wild-type growth was inhibited only above 10 mM, the Δ*tolC* strain was inhibited at concentrations as low as 625 μM (Fig. S5). This indicates that TolC contributes to verapamil tolerance in growth assays, although the underlying mechanism remains unclear, since loss of TolC broadly perturbs efflux, envelope homeostasis, and stress resistance.

Next, we used single-cell BFM recordings to assess whether *tolC* deletion alters the immediate PMF response to verapamil. Motors in the Δ*tolC* background behaved similarly to wild-type: at an intermediate concentration (50 μM), most motors slowed gradually, while a small subpopulation abruptly stopped (see Fig. 2B). We observed both response types in the Δ*tolC* background, similar to wild type. Together, these results suggest that while TolC influences verapamil tolerance at the population level over longer timescales, it does not play a dominant role in shaping the heterogeneous, short-timescale single-cell energetic response.

### 2.3 Ensemble-level swimming speed decreases with verapamil concentration

To complement our single-cell measurements of flagellar motor speed, we performed ensemble-level motility assays using Dynamic Differential Microscopy (DDM) (see Materials and Methods and Supplementary Information Section 4). DDM quantifies the average swimming speed of motile bacteria in bulk liquid and the fraction of motile bacteria by analyzing spatiotemporal intensity fluctuations in image sequences, enabling high-throughput characterization of population-level motility.

As shown in Fig. 2D, the average swimming speed decreased with increasing verapamil concentration. This trend is in line with the speed reduction observed on single motors (Fig. 2A), supporting the conclusion that verapamil impairs PMF-driven motility. However, swimming speed decreased more strongly than motor speed across the same concentration range (Fig. 2D). The fraction of motile cells *α* also decreased with increasing verapamil concentration, indicating a concentration-dependent increase in the number of cells undergoing PMF collapse (Fig. 2D).

### 2.4 Verapamil-induced PMF collapse does not trigger stator unit dissociation

The stator units of the BFM are transmembrane ion channels that couple ion flow across the cytoplasmic membrane to torque generation. In *E. coli*, stator units are proton-driven and can dynamically bind to and unbind from the motor [39], converting PMF into mechanical rotation of the flagellum. Stator recruitment in *E. coli* is mechanosensitive, with the number of engaged units increasing under higher viscous loads, up to approximately a dozen [40–42]. Stator assembly and retention in *E. coli* have also been shown to depend on PMF or on the availability of external coupling ions [43, 44]. Because torque generation in the BFM depends on the engagement of these units, and because classical PMF collapse can alter their occupancy and recovery dynamics, we asked whether the motor arrest induced by verapamil reflects dissociation of stator units from the motor.

To investigate this, we compared the behavior of motors during arrest and recovery following treatment with verapamil or the protonophore CCCP, which collapses the PMF and is known to promote stator unbinding. Consistent with previous reports [34, 43, 45, 46], motors treated with 20 μM CCCP exhibited a gradual stepwise recovery of rotational speed after washout (Fig. 3A), reflecting sequential reincorporation of stator units. In contrast, motors that fully stopped after verapamil treatment rapidly resumed rotation after washout in transitions lasting only a few milliseconds, with no evidence of discrete speed steps (Fig. 3A). We also examined fluctuations of the bead during motor arrest. CCCP-stopped motors showed substantially larger fluctuations than verapamil-stopped motors (Fig. S3). Because these fluctuations report on the mechanical state of the motor-bead system, this observation further supports the conclusion that the arrested state produced by verapamil differs from that induced by a classical protonophore. Together, these results suggest that verapamil-induced motor arrest does not trigger stator dissociation, in contrast to protonophore-induced PMF collapse.

**Figure 3.**
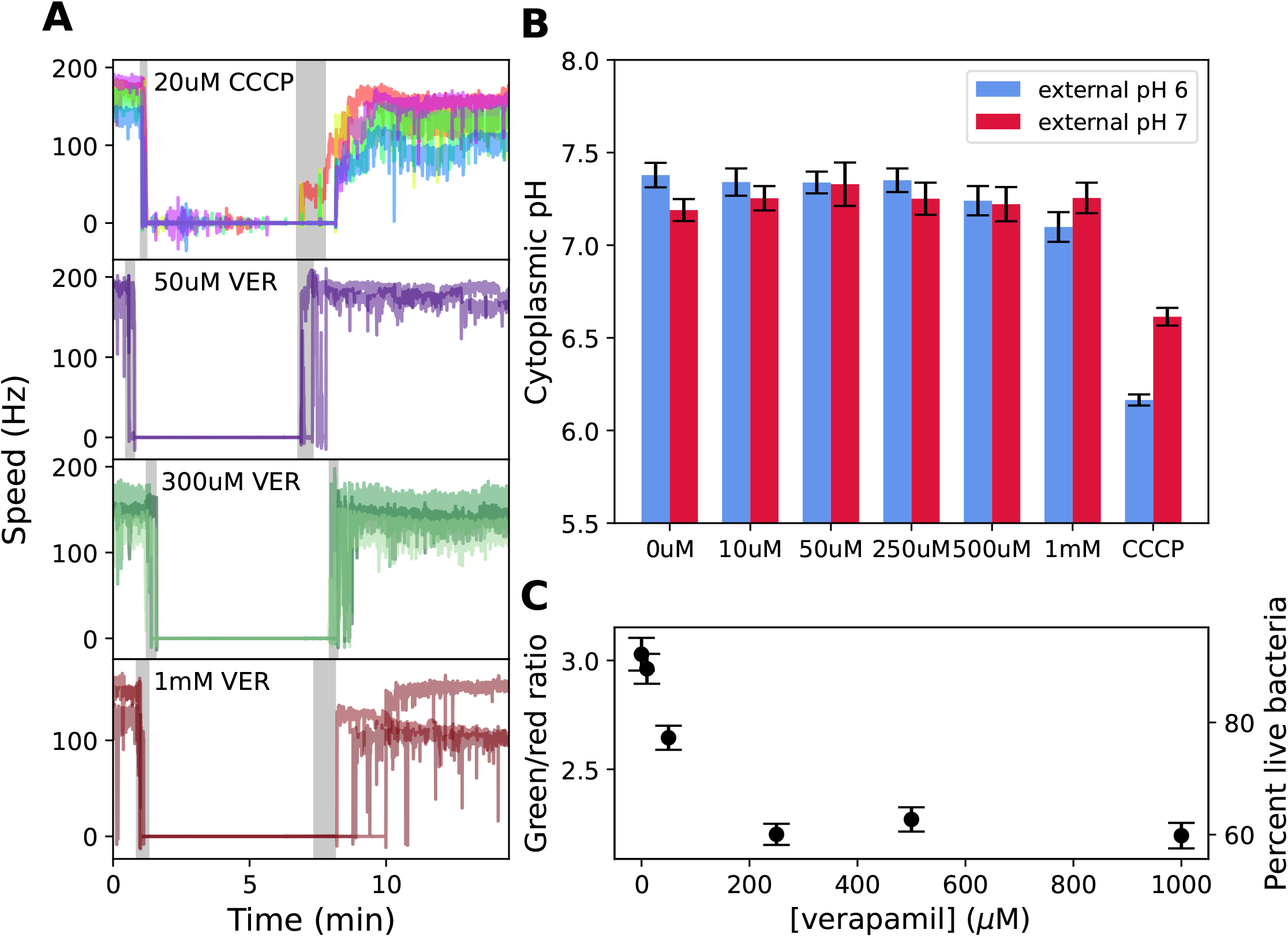
Comparison of recovery dynamics, intracellular pH, and membrane-integrity measurements. (A) Top: motor speed traces following treatment with 20 μM CCCP. After washout, motors recover gradually and in discrete steps, consistent with sequential stator reassembly. Bottom: subset of motors from Fig. 2A that fully stopped during verapamil treatment (concentrations as labeled). After washout, these motors resume rotation rapidly and without detectable stepwise transitions. Grey bars indicate periods during which CCCP or verapamil was present. (B) Intracellular pH measured using pHluorin in the absence or presence of varying concentrations of verapamil or 20 μM CCCP. Measurements were performed in motility buffer (MB) at room temperature during the first 15 min of exposure. Error bars represent standard deviations across three biological replicates. Calibration data are shown in Fig. S6. (C) Green/red fluorescence intensity ratio of bacterial suspensions treated with verapamil and labeled with SYTO9 and propidium iodide (PI). The right y-axis shows the estimated percentage of live cells, calculated from the calibration relating green/red ratio to mixtures of live and heat-killed cells (see Supplementary Information). Each point represents the mean of three independent measurements; error bars were obtained by propagation of uncertainty from the standard deviation of the mean red and green fluorescence intensities (Fig. S8).

### 2.5 Verapamil does not change intracellular pH

Using acridine orange and inverted vesicles, Chen *et al*. reported a concentration-dependent decrease in Δ*pH* in *Mycobacteria bovis BGC* [14]. This effect was also observed in *Mycobacteria smegmantis* by Fountain *et al*., who showed that 100 μM verapamil produced an effect comparable to 10 μM CCCP [13]. However, using radiolabeled benzoic acid uptake, they conversely observed that verapamil had no significant impact on Δ*pH* relative to CCCP. As a potential explanation of such discrepancies, recent work has highlighted that measurements of proton motive force in inverted membrane vesicles and intact cells can yield qualitatively different results for membrane-active compounds, owing to differences in membrane topology, geometry, and charge-state-dependent transport [47]. Given these conflicting results, we sought to directly assess changes in intracellular pH using pHluorin, a noninvasive, rapid, and quantitative pH-sensitive GFP variant [28, 48, 49].

We quantified cytoplasmic pH across a range of external pH values and verapamil concentrations. In parallel, we calibrated pHluorin using known Δ*pH*-collapsing agents (see Materials and Methods and Fig. S6). As shown in Fig. 3B, intracellular pH remained constant within experimental error at both external pH 6 and 7 across the full range of verapamil concentrations tested. We measured an average internal pH of 7.26 *±* 0.10, consistent with bacterial pH homeostasis and previous reports [48, 50]. In contrast, CCCP produced the expected collapse of the ΔpH (Fig. 3B). Although longer incubations (approximately 1.5 h) revealed a modest decrease in intracellular pH under acidic external conditions, no substantial change was observed during the time window relevant to the motor measurements (Fig. S7). These observations suggest that verapamil does not measurably alter intracellular pH over a timescale of approximately one hour.

### 2.6 Verapamil compromises membrane integrity in a dose-dependent manner

To investigate whether verapamil compromises membrane integrity, we used the LIVE/DEAD BacLight bacterial viability kit which employs SYTO9 and propidium iodide (PI) to report on membrane permeability (see Materials and Methods). SYTO9 (green) labels all cells, whereas PI (red) enters only cells with permeabilized membranes and displaces SYTO9 from nucleic acids due to its higher binding affinity. We calibrated the green/red fluorescence ratio using defined mixtures of live and heat-killed cells (see Supplementary Information Section 7). In verapamil-treated cells (Fig. 3C), this ratio decreased with increasing drug concentration, indicating a larger fraction of the population exhibiting PI-permeable membranes.

Further, Minero *et al*. recently reported that verapamil treatment in *Staphylococcus epidermidis* increased SYTO dye uptake, suggesting that SYTO dyes may be substrates of bacterial efflux pumps and that verapamil could inhibit their efflux [51]. Under this interpretation, one would expect increased green fluorescence in the presence of verapamil. In contrast, we observe a decrease in both the green/red fluorescence ratio and the green fluorescence intensity with increasing verapamil concentration (Fig. S9), suggesting that verapamil alters membrane integrity or energetics rather than simply altering dye accumulation.

To assess cultivability, samples from all verapamil concentrations were streaked onto agar plates and incubated at 37 °C for 24 hours. Colonies were observed in all conditions except for the 100% heat-killed control (data not shown), indicating that verapamil-treated cells remain culturable, even when membrane integrity is partially compromised.

Overall, these results are consistent with altered membrane integrity and/or energetics in the presence of verapamil, with the effect saturating at concentrations above 250 μM. However, given the potential influence of membrane potential and efflux activity on dye uptake, these findings should be interpreted with caution. The apparent coexistence of PI uptake with stable intracellular pH may be reconciled if verapamil induces transient and localized membrane defects that permit dye entry but do not generate a sustained proton leak sufficient to overwhelm buffering and active pH homeostasis in respiring cells. Control experiments with CCCP (Fig. S9) revealed that collapsing membrane potential can increase the green/red ratio, demonstrating that this assay is sensitive to membrane energetics in addition to membrane integrity. The fact that verapamil-treated cells remain culturable indicates that membrane damage may be partial or reversible.

### 2.7 Circuit model captures verapamil-induced collapse of membrane potential

To interpret the effect of verapamil on PMF dynamics, we modeled the bacterial membrane using an equivalent electrical circuit, following earlier approaches used to describe ionophore action [28–30, 52–55]. In this lumped-element model, the inner membrane is treated as a capacitor that separates charge between the cytoplasm and the periplasm. We assume the periplasm is in equilibrium with the extracellular environment due to the high permeability of the outer membrane to small ions [56].

The flagellar motor is modeled as a voltmeter, with its rotation speed *ω* linearly proportional to the transmembrane voltage *V*, such that *ω* = *c · V* . Since the external and internal pH are both close to neutral (with *pH*_int_ *≈* 7.2–7.8), the pH gradient contributes only weakly to the PMF. Thus, for our purposes, the PMF can be approximated as purely electrical:

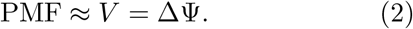

In this model, proton-consuming components such as ATP synthase are represented as a passive load with resistance *R*_*s*,0_, where the subscript *s* denotes a proton sink, while respiration complexes are modeled as a voltage source *V*_*r*_ in series with a resistance *R*_*r*_ that accounts for inefficiencies in catabolism. *V*_*r*_ represents the theoretical maximum membrane potential the cell could achieve in a given environment, relying on a specific internalized carbon source.

To account for the verapamil-induced collapse of the PMF, we include an additional leak pathway across the membrane: a switch-controlled resistor *R*_*v*_ connected in parallel to *R*_*s*,0_. This branch is activated (i.e., the switch closes) when verapamil is introduced, modeling increased membrane conductance. This phenomenological term does not assume a specific microscopic mechanism, but captures any process that alters the effective electrical properties of the membrane, including increased ion permeability, transient defects, or changes in membrane electrostatics. The total sink resistance then becomes:

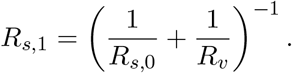

The solution for the time-dependent transmembrane voltage is:

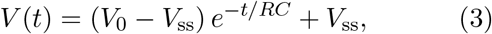

where *C* is the membrane capacitance, *V*_0_ is the initial membrane potential, and the effective resistance is *R* = (1*/R*_*r*_ + 1*/R*_*s*_)^−1^. The steady-state membrane potential after verapamil addition is:

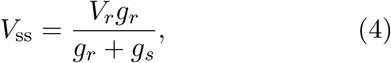

where *g*_*r*_ = 1*/R*_*r*_ and *g*_*s*_ = 1*/R*_*s*_ are the conductances of the respiration source and the proton sinks, respectively. In the limit of very high effective membrane conductance (large *g*_*s*_), the steady-state voltage approaches zero, reflecting a full PMF collapse.

If we assume the verapamil-induced conductance scales linearly with its concentration, i.e., *g*_*v*_ = *γ*[Ver], then the normalized steady-state voltage after verapamil addition becomes:

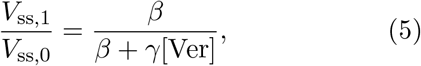

where *β* = *g*_*s*,0_+*g*_*r*_, and *V*_ss,0_ is the membrane potential prior to verapamil addition (switch open), while *V*_ss,1_ denotes the membrane potential after verapamil addition (switch closed).

Fig 2C shows the average motor speed traces (proportional to the membrane voltage) for cells that retained measurable rotation following verapamil treatment across several concentrations. These traces were fitted with a single exponential decay function. In Fig. 2D, we plot the average steady-state motor speed (from the final seconds before washout) at each verapamil concentration. These normalized values were then fit to the hyperbolic function in Eq. 5, allowing us to extract the empirical parameters *γ* and *β*, which reflect the drug-induced membrane permeability and the baseline proton conductance, respectively.

## 3 Discussion

In this work we aimed to reconcile the seemingly contradictory views on verapamil’s role in potentiating antibiotics against Mtb. Verapamil, a calcium channel blocker and efflux inhibitor, has been proposed to act primarily by inhibiting drug efflux pumps, thereby enhancing the intracellular accumulation of antibiotics. However, other studies have suggested that it may also act by disrupting membrane energetics. By combining single-cell electrophysiological measurements of bacterial flagellar motor speed, population-level growth and swimming assays, and mutant analysis, we explored how and when verapamil perturbs bacterial physiology.

Our motor speed measurements demonstrate that verapamil causes a concentration-dependent reduction of the PMF in *E. coli*, and that this response is largely reversible upon washout, especially at low to moderate concentrations. Our DDM measurements confirm these observations, revealing a similar concentration-dependent decrease in swimming speed at the population level (Fig. 2D). In addition, DDM quantifies the population motile fraction *α* and the Brownian diffusion coefficient *D* (Fig. 2D and Fig. S4) [57]. We note that the magnitude of the speed decrease obtained by DDM is greater than that observed in the single motor measurements (Fig. 2D). Part of this discrepancy possibly arises because effective swimming requires stable flagellar bundle formation, which depends on the coordinated rotation of multiple filaments; even small reductions in motor speed can disrupt bundling and disproportionally reduce propulsion [58, 59]. Moreover, when *α* becomes small, the fitted parameters become partially correlated, and cross-talk between them can bias the fitted speed downward [60–62]. Finally, the two assays probe slightly different viscous load regimes, so the relative fractional change in speed may be different in the two experiments.

The verapamil-induced speed reduction observed at the single motor level was characterized by a bimodal behavior. While bead assays do not allow precise quantification of the relative frequencies of these two behaviors, we see that the majority of motors slow gradually, while a subpopulation abruptly stop. To test whether this heterogeneity was driven by efflux pump activity, we examined a Δ*tolC* mutant lacking the outer membrane channel required for AcrAB-TolC-mediated export. In growth assays, this strain exhibited increased sensitivity to verapamil, with growth inhibition occurring at concentrations above approximately 625 μM, compared to inhibition above 10 mM in the wild type. For comparison, *M. tuberculosis* shows a verapamil minimum inhibitory concentration (MIC) of 0.5–1.1 mM [63], suggesting that *E. coli* is intrinsically about tenfold more tolerant. Disruption of AcrAB-TolC therefore brings *E. coli* sensitivity closer to that of *M. tuberculosis*, consistent with prior work showing that loss of TolC increases verapamil susceptibility by an order of magnitude [22]. Despite this increased sensitivity, single-cell measurements showed that Δ*tolC* cells displayed the same bimodal behavior as wild type. In contrast to protonophore-induced bimodality, which can arise from an efflux-mediated positive feedback loop [32], our results suggest that TolC-mediated efflux does not explain the cell-to-cell heterogeneity in verapamil-induced PMF disruption. Instead, this heterogeneity may reflect pre-existing physiological differences between cells, including variations in metabolic state and baseline PMF, membrane composition and electrostatics, efflux capacity, or variability in the effective membrane-associated concentration of verapamil across cells.

Fountain *et al*. [13] recently showed that verapamil and its metabolite norverapamil inhibit the MmpS5–MmpL5 efflux pump in *Mycobacterium tuberculosis*, enhancing the activity of bedaquiline. Their work supports an efflux inhibition model and argues against membrane depolarization as a physiologically relevant mechanism, reporting no significant verapamil-induced changes in Δ*ψ* or ΔpH in *M. smegmatis*. Our findings suggest an additional mechanism not captured by this framework: because many efflux pumps, including MmpL5, are powered by the PMF [64], even transient or partial energetic disruption would be expected to impair their function. Thus, part of the observed efflux inhibition may reflect secondary consequences of altered membrane energetics, rather than, or in addition to, direct binding to the pump.

To quantify the energetic effects, we modeled the membrane as a capacitor with resistive leak channels. This phenomenological circuit model captures the observed hyperbolic dependence of motor speed on verapamil concentration without assuming a specific molecular mechanism and yields membrane resistance values consistent with prior literature [30, 53]. Our LIVE/DEAD BacLight staining results further support the idea that verapamil compromises membrane integrity, though interpretation is complicated by dye sensitivity to both membrane potential and efflux [65, 66].

In proton-driven *E. coli*, the collapse of the PMF by the protonophore CCCP leads to motor recovery with discrete speed steps, commonly interpreted as stator dissociation followed by sequential reassembly [34, 43, 45]. Similar stepwise “resurrection” dynamics have been observed in electrorotation experiments, where transient reduction of motor torque drives stator dissociation, followed by discrete reassembly on minute timescales upon restoration of load [67, 68]. Fluorescence-based measurements, however, yield mixed results: several studies report stator unit disengagement following PMF collapse [43, 69], while others observed preserved stator localization even in the presence of CCCP [44]. In contrast, following verapamil treatment we observe near-instant restoration of pre-treatment motor speed after washout, without detectable resurrection steps. This behavior is consistent with stator units remaining engaged throughout verapamil-induced motor arrest and is similar to recent observations under hyperosmotic stress [31]. Consistent with this interpretation, motors arrested by verapamil exhibited markedly smaller bead fluctuations than CCCP-arrested motors, indicating that the mechanical state of the motor remains distinct from that produced by classical protonophore treatment (Fig. S3). Together, these observations point to a mechanistic distinction between verapamil-induced motor arrest and the PMF collapse caused by classical protonophores.

Prior work in proton-driven motors of *E. coli* and *Salmonella enterica* has shown that stator assembly and retention depend primarily on external proton availability, and that lowering external pH can restore stator occupancy even when the membrane potential is impaired [44, 69]. This suggests that preservation of a transmembrane proton gradient can maintain stator engagement even if Δ*ψ* is collapsed. We thus hypothesize that verapamil does not act as a classical protonophore, but instead may primarily dissipate Δ*ψ*, while leaving ΔpH at least partially intact, consistent with our pHluorin measurements showing stable intracellular pH in *E. coli* over the timescales probed. This is supported by measurements showing that verapamil decreases Δ*ψ* in *E. coli* at concentrations above *∼* 0.5 mM [22]. It also aligns with prior biophysical studies showing that verapamil alters membrane dipole potential and affects membrane permeability without acting as a direct proton shuttle, as well as with structural studies showing perturbations to membrane organization that may influence ion transport [20, 21, 23, 25]. Similar “electrostatic smoothing” effects have been reported for other amphiphilic compounds like local anesthetics, which can reduce the membrane potential sensed by proteins without extensive proton leak [70]. Finally, this view is supported by measurements in *M. tuberculosis* showing rapid depolarization but only partial dissipation of the proton gradient in the presence of verapamil [14]. Proton-coupled secondary transporters are also sensitive to membrane electrostatics and could, in principle, be influenced by verapamil’s membrane-partitioning behavior.

Overall, our results support a model in which verapamil accumulates within the inner membrane lipid bilayer and causes a concentration-dependent, reversible perturbation of membrane energetics, likely through preferential dissipation of the electrical component of the PMF. While our higher verapamil concentrations extend beyond the range commonly used in physiologically relevant assays, the verapamil-induced decrease in PMF appears to follow a continuum across concentrations. On short timescales, this membrane-associated effect produces heterogeneous energetic disruption while leaving the flagellar motor and its stator units intact, enabling rapid functional recovery. As these measurements probe relatively short timescales (minutes), they likely capture early, dynamic consequences of membrane perturbation, whereas longer exposures may lead to more pronounced effects. On longer timescales, continued accumulation of verapamil in the membrane may contribute to growth inhibition and antibiotic potentiation.

Our findings do not rule out direct efflux pump inhibition, but they suggest that energetic disruption may also contribute by reducing the PMF available to power efflux systems. In addition, verapamil has also been reported to downregulate efflux pump expression and reduce resistance emergence over longer timescales [71]. While our measurements capture rapid, reversible perturbations of membrane energetics on the order of minutes, changes in gene expression and resistance modulation emerge over hours to days, suggesting that multiple mechanisms may act sequentially or in parallel. Rather than acting exclusively as either an efflux pump inhibitor or a membrane disruptor, verapamil likely exerts dual-action effects that sensitize bacteria to antibiotics without killing them outright. This has important implications for TB therapy, especially under host conditions where membrane energetics and efflux capacity are dynamically regulated and may jointly influence antibiotic susceptibility.

## 4 Materials and Methods

### Bacterial strain and culture

For bead assay experiments, we used *E. coli* strain MT03, a derivative of RP437 in which *cheY* is deleted (producing a counterclockwise-only phenotype) and *fliC* is replaced by the *fliC*^*st*^ allele encoding “sticky” flagellar filaments that facilitate bead attachment. For DDM experiments, we constructed an MG1655 Δ*cheY* strain, a smooth swimming mutant. To probe the role of TolC-mediated efflux, we generated an MT03 Δ*tolC* strain. Gene deletions were introduced by *λ*-Red recombineering, replacing *cheY* and *tolC* with a spectinomycin-resistance cassette [72]. For pH measurements, we used strain MT03 harboring plasmid pYVM063, which expresses ratiometric pHluorin under the arabinose-inducible promoter *P*_*BAD*_ [73] (a kind gift from the lab of Tohru Minamino).

Unless otherwise noted, cultures were initiated by inoculating 100 μL of frozen stock (stationary-phase culture stored in 25% glycerol at −80 °C) into 5 mL of Lysogeny Broth (LB; 10 g L^−1^ bactotryptone, 5 g L^−1^ yeast extract, 10 g L^−1^ NaCl). Cultures were grown aerobically at 35 °C in a shaking incubator at 200 rpm for approximately 4 hours. Cells were harvested at mid-to late-log phase, corresponding to an OD_600_ of 0.6–0.8.

### Bead assay

Flagella were mechanically sheared by passing 1 mL of cell culture back and forth between two syringes connected by 21-gauge needles and a narrow connecting tube, as described previously [74]. Cells were then centrifuged at 3000 rpm for 2 min. After discarding the supernatant, the cell pellet was re-suspended in 300 μL of motility buffer (MB; 10 mM potassium phosphate, 0.1 mM EDTA, 10 mM lactic acid, pH 7.0).

Cells were introduced into custom tunnel slides prepared from two Menzel-Gläser #1.5 coverslips separated by a layer of parafilm in which a central channel had been cut. Two small holes at the extremities of the top coverslip, made using a laser cutter, allowed fluid exchange [33]. Prior to use, tunnel slides were cleaned by flushing sequentially with ethanol and sterile water. To adhere the cells, 100 μL of poly-L-lysine solution (Sigma-Aldrich, P4707) was flushed into the tunnel slide and incubated for 2 minutes, then washed out with 200 μL of MB. Next, 200 μL of the prepared cell suspension was introduced into the tunnel slide, and cells were allowed to settle and adhere for 10 minutes. Non-adherent cells were then washed out with MB.

A 300 μL aliquot of a 1:300 dilution of 600 nm diameter polystyrene beads (Sigma-Aldrich, LB6) in MB was flushed through the tunnel slide. Beads were allowed to bind spontaneously to the truncated “sticky” filaments, and after 10 min unbound beads were removed by flushing with MB. Motor rotation was recorded by tracking bead motion on a custom-built bright-field microscope with 660 nm LED illumination (Thorlabs M660L3), a Nikon 100× 1.45 NA oil-immersion objective, and a high-speed CMOS camera (Optronics CL600x2/M). Rotating beads were imaged at 1 kHz.

Rotating beads were recorded for approximately 1 min to establish baseline motor speed. Verapamil solutions (200 μL in MB at the desired concentration) were then introduced into the tunnel slide, followed by a 5–10 min incubation and washout with 800 μL fresh MB. CCCP experiments were performed similarly, with 20 μM CCCP introduced after baseline recording, followed by the same incubation and washout procedure.

### Dynamic Differential Microscopy

An overnight culture of MG1655 Δ*cheY* was prepared using the growth conditions described above. The following day, a fresh culture was started by diluting 200 μL of the overnight culture into 10 mL of LB and incubating under the same conditions until the OD_600_ reached *∼*0.8. All cultures contained μL of a 100 mg/mL spectinomycin stock solution. Cells were harvested and washed three times in motility buffer (MB) by centrifugation at 4000 rpm for 2 min. After the final wash, the pellet was resuspended in MB and diluted to OD_600_ *∼* 0.3 for DDM imaging.

Immediately prior to imaging, verapamil was added to the bacterial suspension to the desired final concentration, and 100 μL of the mixture was loaded into a tunnel slide constructed from a 24 *×* 60 mm *×* coverslip (bottom) and a 22 × 22 mm coverslip (top), separated by a 265 μm layer of double-sided tape (Menzel-Gläser #1.5). The ends of the tunnel were sealed with vacuum grease to prevent evaporation. DDM measurements were made without verapamil at both the beginning and end of the experiment to confirm that motility remained stable over the measurement period; the entire procedure, from the first control acquisition to the final one, was completed within one hour. We used an inverted microscope (Nikon, eclipse Ti) and a high-speed camera (Photron Fastcam Nova S16) with a fast drive. Bacteria were imaged with a 10x phase-contrast objective with a 512 x 512 pixel field of view for 3 min at 250 fps.

Swimming speeds were quantified by Dynamic Differential Microscopy (DDM) using established methods [60, 75]. Briefly, image time series were analyzed to compute the differential intensity correlation function, from which the intermediate scattering function (ISF) was obtained at each wave vector *q*. ISFs were fitted using a standard model for smooth-swimming *E. coli*, consisting of a mixture of diffusive and ballistic (swimming) components. The fitted parameters were the motile fraction *α* and the mean swimming speed 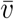, with the shape of the speed distribution fixed (see SI, Section 4 for details of the ISF model, the Schulz distribution, and the constrained fitting procedure).

### pHluorin assay

MT03 pHluorin cells were grown in 96-well deep plates containing 0.5 mL LB supplemented with chloramphenicol (25 μg*/*mL), 0.01 % arabinose, and a 1:50 dilution of an overnight culture. The plates were covered with a breathable seal (AeraSeal film, Sigma-Aldrich, A9224) and incubated under the previously described growth conditions. After 4 h, cells were centrifuged at 2000 g for 10 min. The pellet was resuspended in MB, and this wash step was repeated once. Cells were then incubated in MB for 30 min, followed by a third centrifugation and re-suspension via a 96-well pipetting robot in one of the following buffers: MB supplemented with 20 μM CCCP (pH adjusted after CCCP addition; final Ph range 5.5–9), or 40 mM PBMH (potassium benzoate and methylamine hydrochloride; pH adjusted after buffer preparation; final pH range 5.5–9). Aliquots of 150 μL of cells in each buffer condition were transferred into a black 96-well microplate (Greiner, 655090). Fluorescence was measured using a Synergy HTX microplate reader (BioTek Instruments) with excitation at 395 nm and 475 nm, and emission detected at 515 nm. Measurements were taken every 5 min at 25 °C.

### LIVE/DEAD fluorescence assay

An overnight culture of MT03 was prepared using the growth conditions described above. The next day, a fresh culture was initiated by diluting 1 mL of the overnight culture into 50 mL of LB and grown to mid-log phase. Cells were harvested by centrifugation at 4000×*g* for 10 min and washed twice with motility buffer (MB). A heat-killed control (100% dead) was generated by incubating an aliquot at 95 °C for 15 min in a water bath.

Bacterial suspensions containing different verapamil concentrations were prepared in MB to a final volume of 100 μL. An equal volume (100 μL) of the SYTO9/propidium iodide mixture from the LIVE/DEAD BacLight kit (L7012, ThermoFisher) was added according to the manufacturer’s protocol. Calibration standards consisting of defined mixtures of live and heat-killed cells (0–100%) were prepared in parallel.

After a 15 min incubation at room temperature, each sample was transferred to a black 96-well microplate (Greiner, 655090). Fluorescence was measured on a Synergy HTX microplate reader (BioTek Instruments) using 485 nm excitation with dual-emission detection at 530 nm (SYTO9) and 630 nm (propidium iodide). OD_600_ was recorded to normalize for cell density. The green/red fluorescence ratios were compared to the calibration curve to estimate the percentage of viable cells. To assess cultivability, 10 μL of each suspension was streaked onto LB agar plates and incubated at 37 °C for 24 h.

## Acknowledgments

We thank Mark Troll for inspiring this work and for invaluable insights and feedback throughout. We are grateful to Alexandra Lake and Greg Cook for careful reading of the manuscript and helpful comments. We thank Pauline Mayonove and Jérôme Bonnet for advice, technical support, and access to equipment for plate reader measurements. We also thank the laboratory of Tohru Minamino for providing the pHluorin plasmid. A.L.N., A.B.B., M.M., N.M., and P.L.V. were supported by the Impulscience project PHYBION of the Bettencourt Schueller Foundation. A.L.N., A.A., and P.L.V. were further supported by the ANR PHYBABIFO project (ANR-22-CE30-0034). A.L.N. and A.B.B. were also supported by the ANR PHYBION project (ANR-23-ERCB-0005-01). F.P. and A.A. were supported by the ANR BaElMec project (ANR-23-CE30-0010). The CBS is a member of France-BioImaging (FBI) and the French Infrastructure for Integrated Structural Biology (FRISBI), two national infrastructures supported by the French National Research Agency (ANR-10-INBS-04-01 and ANR-10-INBS-05, respectively).

## References

1. World Health Organization. Global Tuberculosis Report 2024 https://www.who.int/publications/i/item/9789240093929 (World Health Organization, Geneva, 2024).

2. Mitini-Nkhoma, S. C., Fernando, N., Ishaka, G. K. D., Handunnetti, S. M. & Pathirana, S. L. Ion transport modulators as antimycobacterial agents. Tuberculosis Research and Treatment 2020, e3767915 (2020).

3. Omollo, C. et al. Developing synergistic drug combinations to restore antibiotic sensitivity in drug-resistant Mycobacterium tuberculosis. Antimicrobial Agents and Chemotherapy 65, e02554.#x2013;20, AAC.02554–20 (2023).

4. McGoon, M. D., Vlietstra, R. E., Holmes, D. R. & Osborn, J. E. The clinical use of verapamil. Mayo Clinic Proceedings 57, 495–510 (1982).

5. Gupta, S. et al. Acceleration of tuberculosis treatment by adjunctive therapy with Verapamil as an efflux inhibitor. American Journal of Respiratory and Critical Care Medicine 188, 600–607 (2013).

6. Gupta, S. et al. Efflux inhibition with Verapamil potentiates bedaquiline in Mycobacterium tuberculosis. Antimicrobial Agents and Chemotherapy 58, 574–576 (2014).

7. Adams, K. N. et al. Drug tolerance in replicating mycobacteria mediated by a macrophage-induced efflux mechanism. Cell 145, 39–53 (2011).

8. Pieterman, E. D. et al. Assessment of the additional value of verapamil to a moxifloxacin and Linezolid Combination regimen in a murine tuberculosis model. Antimicrobial Agents and Chemotherapy 62, e01354–18 (2018).

9. Adams, K. N., Szumowski, J. D. & Ramakrishnan, L. Verapamil, and its metabolite norverapamil, inhibit macrophage-induced, bacterial efflux pump-mediated tolerance to multiple anti-tubercular drugs. The Journal of Infectious Diseases 210, 456–466 (2014).

10. Xu, J. et al. Verapamil increases the bioavailability and efficacy of bedaquiline but not clofazimine in a murine model of tuberculosis. Antimicrobial Agents and Chemotherapy 62, e01692–17 (2018).

11. Lake, M. A. et al. The human proton pump inhibitors inhibit Mycobacterium tuberculosis rifampicin efflux and macrophage-induced rifampicin tolerance. Proceedings of the National Academy of Sciences 120, e2215512120 (2023).

12. Tang, L. et al. Structural basis for inhibition of a voltage-gated Ca2+ channel by Ca2+ antagonist drugs. Nature 537. issn: 1476-4687 (2016).

13. Ramakrishnan, G., Ravindran, M., Radhakrishnan, A., DeMajistre, R. & Manjunatha, U. Verapamil and its metabolite norverapamil inhibit the mycobacterium tuberculosis MmpS5–MmpL5 Efflux Pump. Antimicrobial Agents and Chemotherapy 69, e01173–23 (2025).

14. Chen, C. et al. Verapamil targets membrane energetics in mycobacterium tuberculosis. Antimicrobial Agents and Chemotherapy 62, e02107–17 (2018).

15. Te Winkel, J. D., Gray, D. A., Seistrup, K. H., Hamoen, L. W. & Strahl, H. Analysis of antimicrobial-triggered membrane depolarization using voltage sensitive dyes. Frontiers in cell and developmental biology 4, 29 (2016).

16. Valderrama, K. et al. Pyrrolomycins are potent natural protonophores. Antimicrobial agents and chemotherapy 63, 10–1128 (2019).

17. Mancini, L. et al. A general workflow for characterization of nernstian dyes and their effects on bacterial physiology. Biophysical Journal 118, 4–14 (2020).

18. Buttress, J. A. et al. A guide for membrane potential measurements in Gram-negative bacteria using voltage-sensitive dyes. Microbiology 168, 001227 (2022).

19. Lo, W.-C., Krasnopeeva, E. & Pilizota, T. Bacterial electrophysiology. Annual Review of Biophysics 53 (2024).

20. Meier, M., Blatter, X. L., Seelig, A. & Seelig, J. Interaction of verapamil with lipid membranes and P-glycoprotein: connecting thermodynamics and membrane structure with functional activity. Biophysical Journal 91, 2943–2955 (2006).

21. Pohl, E. E., Krylov, A. V., Block, M. & Pohl, P. Changes of the membrane potential profile induced by verapamil and propranolol. Biochimica Et Biophysica Acta 1373, 170–178 (1998).

22. Andersen, C. L., Holland, I. B. & Jacq, A. Ve-rapamil, a Ca2+ channel inhibitor acts as a local anesthetic and induces the sigma E dependent extra-cytoplasmic stress response in E. coli. en. Biochimica et Biophysica Acta (BBA) - Biomembranes 1758, 1587–1595 (2006).

23. Phua, Z. Y. et al. A potent phenylalkylamine disrupts mycobacterial membrane bioenergetics and augments bactericidal activity of bedaquiline. iScience (2025).

24. Spycher, S., Smejtek, P., Netzeva, T. I. & Escher, B. I. Toward a class-independent quantitative structure-activity relationship model for uncouplers of oxidative phosphorylation. Chemical Research in Toxicology 21, 911–927 (2008).

25. Suwalsky, M. et al. Structural effects of verapamil on cell membranes and molecular models. Journal of the Chilean Chemical Society 55, 1– 4 (2010).

26. Gabel, C. V. & Berg, H. C. The speed of the flagellar rotary motor of Escherichia coli varies linearly with protonmotive force. Proceedings of the National Academy of Sciences 100, 8748– 8751 (2003).

27. Krasnopeeva, E. et al. Nonlinear dependency of the bacterial flagellar motor speed on proton motive force and its consequences for swimming. bioRxiv (2025).

28. Krasnopeeva, E., Lo, C.-J. & Pilizota, T. Single-cell bacterial electrophysiology reveals mechanisms of stress-induced damage. Bio-physical Journal 116, 2390–2399 (2019).

29. Krasnopeeva, E., Barboza-Perez, U. E., Rosko, J., Pilizota, T. & Lo, C.-J. Bacterial flagellar motor as a multimodal biosensor. Correlative approaches in single-molecule biophysics: a review of the progress in methods and applications 193, 5–15 (2021).

30. Biquet-Bisquert, A. et al. Spatiotemporal dynamics of the proton motive force on single bacterial cells. eng. Science Advances 10, eadl5849 (2024)

31. Meneses, L., Dudebout, E. M., Belser, S., Yang, J. & Wadhwa, N. Osmotic stress triggers fast and reversible PMF collapse in Escherichia coli. bioRxiv (2025).

32. Le, D., Krasnopeeva, E., Sinjab, F., Pilizota, T. & Kim, M. Active efflux leads to heterogeneous dissipation of proton motive force by protonophores in bacteria. MBio 12, 10–1128 (2021).

33. Hoffmann, W. H., Biquet-Bisquert, A., Pedaci, F. & Nord, A. L. in Molecular Motors: Methods and Protocols 43–64 (Springer, 2024).

34. Perez-Carrasco, R. et al. Relaxation time asymmetry in stator dynamics of the bacterial flagellar motor. Science Advances 8, eabl8112 (2022).

35. Krulwich, T. A., Sachs, G. & Padan, E. Molecular aspects of bacterial pH sensing and homeostasis. Nature Reviews Microbiology 9, 330– 343 (2011).

36. Koronakis, V., Eswaran, J. & Hughes, C. Structure and function of TolC: the bacterial exit duct for proteins and drugs. Annual review of biochemistry 73, 467–489 (2004).

37. Li, X.-Z., Plésiat, P. & Nikaido, H. The challenge of efflux-mediated antibiotic resistance in Gram-negative bacteria. Clinical microbiology reviews 28, 337–418 (2015).

38. Augustus, A. M., Celaya, T., Husain, F., Humbard, M. & Misra, R. Antibiotic-sensitive TolC mutants and their suppressors. Journal of bacteriology 186, 1851–1860 (2004).

39. Leake, M. C. et al. Stoichiometry and turnover in single, functioning membrane protein complexes. Nature 443, 355–358 (2006).

40. Lele, P. P., Hosu, B. G. & Berg, H. C. Dynamics of mechanosensing in the bacterial flagellar motor. Proceedings of the National Academy of Sciences 110, 11839–11844 (2013).

41. . Nord, A. L. et al. Catch bond drives stator mechanosensitivity in the bacterial flagellar motor. Proceedings of the National Academy of Sciences 114, 12952–12957 (2017).

42. Tipping, M. J., Delalez, N. J., Lim, R., Berry, R. M. & Armitage, J. P. Load-dependent assembly of the bacterial flagellar motor. MBio 4, 10–1128 (2013).

43. Tipping, M. J., Steel, B. C., Delalez, N. J., Berry, R. M. & Armitage, J. P. Quantification of flagellar motor stator dynamics through in vivo proton-motive force control. Molecular microbiology 87, 338–347 (2013).

44. . Suzuki, Y. et al. Effect of the MotA(M206I) mutation on torque generation and stator assembly in the Salmonella H+-driven flagellar motor. Journal of Bacteriology 201, 10.1128/jb.00727–18 (2019).

45. Blair, D. F. & Berg, H. C. Restoration of torque in defective flagellar motors. Science 242, 1678–1681 (1988).

46. Nord, A. L. et al. Dynamic stiffening of the flagellar hook. Nature Communications 13, 2925 (2022).

47. . Harrison, S. H. et al. Remission spectroscopy resolves the mechanism of action of bedaquiline within living mycobacteria. Nature Communications 16, 11018 (2025).

48. Martinez, K. A. et al. Cytoplasmic pH response to acid stress in individual cells of Escherichia coli and Bacillus subtilis observed by fluorescence ratio imaging microscopy. Applied and environmental microbiology 78, 3706–3714 (2012).

49. Wang, Y.-K. et al. Comparison of Escherichia coli surface attachment methods for single-cell microscopy. Scientific reports 9, 19418 (2019).

50. Terradot, G., Krasnopeeva, E., Swain, P. S. & Pilizota, T. Escherichia coli maintains pH via the membrane potential. PRX Life 2, 043015 (2024).

51. Minero, G. A. S., Larsen, P. B., Hoppe, M. E. & Meyer, R. L. Bacterial efflux pumps excrete SYTO− dyes and lead to false-negative staining results. Analyst 149, 2232–2235 (2024).

52. Van Rotterdam, B. J., Crielaard, W., Van Stokkum, I. H., Hellingwerf, K. J. & Westerhoff, H. V. Simplicity in complexity: the photosynthetic reaction center performs as a simple 0.2 V battery. FEBS letters 510, 105–107 (2002).

53. Walter, J. M., Greenfield, D., Bustamante, C. & Liphardt, J. Light-powering Escherichia coli with proteorhodopsin. Proceedings of the National Academy of Sciences 104, 2408–2412 (2007).

54. Arlt, J., Martinez, V. A., Dawson, A., Pilizota, T. & Poon, W. C. Painting with light-powered bacteria. Nature communications 9, 768 (2018).

55. Frangipane, G. et al. Dynamic density shaping of photokinetic E. coli. Elife 7, e36608 (2018).

56. Sun, J., Rutherford, S. T., Silhavy, T. J. & Huang, K. C. Physical properties of the bacterial outer membrane. Nature Reviews Microbiology 20, 236–248 (2022).

57. Martinez, V. & Asally, M. Differential dynamic microscopy for rapid characterization of bacterial motility and viability. Journal of Bacteriological Methods 193, 106358 (2022).

58. Turner, L., Ryu, W. S. & Berg, H. C. Real-time imaging of fluorescent flagellar filaments. Journal of Bacteriology 182, 2793–2801 (2000).

59. Darnton, N. C., Turner, L., Rojevsky, S. & Berg, H. C. On torque and tumbling in swimming Escherichia coli. Journal of Bacteriology 189, 1756–1764 (2007).

60. Wilson, L. G. et al. Differential dynamic microscopy of bacterial motility. Physical review letters 106, 018101 (2011).

61. Martinez, V. et al. Differential dynamic microscopy: a high-throughput method for characterizing the motility of microorganisms. Biophysical journal 103, 1637–47 (Oct. 2012).

62. Germain, D., Leocmach, M. & Gibaud, T. Differential dynamic microscopy to characterize Brownian motion and bacteria motility. American Journal of Physics 84, 202–210 (2016).

63. Machado do Nascimento, A. A. et al. Verapamil Modulates Activity of Antimicrobials Against Rapidly Growing Mycobacteria. Microbial Drug Resistance 31, 162–167 (2025).

64. Poole, K. Efflux pumps as antimicrobial resistance mechanisms. Frontiers in Bioscience 12, 2414–2425 (2007).

65. Berney, M., Hammes, F., Bosshard, F., Weilenmann, H.-U. & Egli, T. Assessment and interpretation of bacterial viability by using the LIVE/DEAD BacLight kit in combination with flow cytometry. Applied and Environmental Microbiology 73, 3283–3290 (2007).

66. Breeuwer, P. & Abee, T. A novel method for monitoring microbial cell membrane integrity using SYTO9 and propidium iodide. Letters in Applied Microbiology 21, 255–259 (1995).

67. Wadhwa, N., Phillips, R. & Berg, H. C. Torque-dependent remodeling of the bacterial flagellar motor. Proceedings of the National Academy of Sciences 116, 11764–11769 (2019).

68. Wadhwa, N., Tu, Y. & Berg, H. C. Mechanosensitive remodeling of the bacterial flagellar motor is independent of direction of rotation. Proceedings of the National Academy of Sciences 118, e2024608118 (2021).

69. Fung, D. C. & Berg, H. C. Powering the flagellar motor of Escherichia coli with an external voltage source. Nature 375, 809–812 (1995).

70. Högberg, C.-J. & Lyubartsev, A. P. Effect of local anesthetic lidocaine on electrostatic properties of a lipid bilayer. Biophysical Journal 94, 525–531 (2008).

71. Liu, K. et al. Verapamil suppresses the development of resistance against anti-tuberculosis drugs in mycobacteria. International Journal of Molecular Sciences 26, 11124 (2025).

72. Datsenko, K. A. & Wanner, B. L. One-step inactivation of chromosomal genes in ¡i¿Escherichia coli¡/i¿ K-12 using PCR products. Proceedings of the National Academy of Sciences 97, 6640–6645 (2000).

73. Morimoto, Y., Kami-Ike, N., Namba, K. & Minamino, T. Determination of local pH differences within living Salmonella cells by high-resolution pH imaging based on pH-sensitive GFP derivative, pHluorin(M153R). Bio Protov 7, e2529 (Sept. 2017).

74. Gabel, C. V. & Berg, H. C. The speed of the flagellar rotary motor of Escherichia coli varies linearly with protonmotive force. Proceedings of the National Academy of Sciences 100, 8748–8751 (2003).

75. Martinez, V. A. et al. Differential dynamic microscopy: a high-throughput method for characterizing the motility of microorganisms. Biophysical journal 103, 1637–1647 (2012).

