## Supplementary material for "Single-Cell Electrophysiology Reveals Verapamil’s Disruption of Bacterial Membrane Energetics": SI

#### Contents

**1 Simultaneous BFM measurements reveal heterogeneity in PMF response 2**

**2 Speed recovery 2**

**3 Fluctuations during motor stops 2**

**4 Differential Dynamic Microscopy 3**

**5 Antimicrobial susceptibility assay 6**

**6 Internal pH measurements 7**

**7 LIVE/DEAD fluorescence for viability estimation 7**

**8 Steady-state motor speeds in $\Delta$*tolC* and wild-type strains 9**

### 1 Simultaneous BFM measurements reveal heterogeneity in PMF response

To illustrate the cell-to-cell variability observed in response to verapamil, Fig. S1 shows the rotational speeds of multiple bacterial flagellar motors recorded simultaneously in the same field of view. These data highlight the heterogeneous motor responses described in the main text.

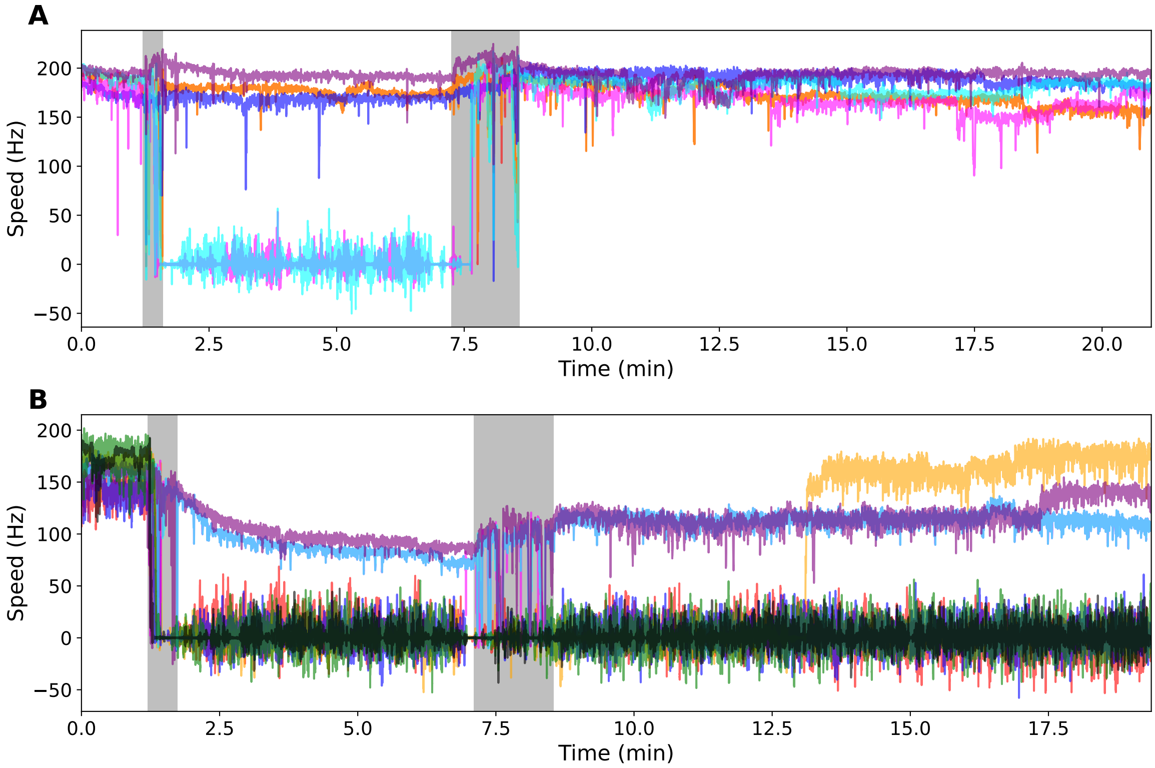


**Figure S1:** Simultaneous measurements of BFM speed from multiple cells in the same field of view illustrate the heterogeneous effect of verapamil on the PMF. (A) At 50 µM verapamil, five motors were tracked simultaneously; some motors exhibit a gradual decline in speed, while others stop abruptly. (B) At 500 µM verapamil, eight motors were recorded in parallel, showing a similarly heterogeneous response.

### 2 Speed recovery

Fig. S2 shows motor speed following verapamil washout. For concentrations up to 300 µM, motor speed generally returned to pre-treatment levels, whereas higher concentrations frequently resulted in partial recovery.

### 3 Fluctuations during motor stops

To further compare the physiological states induced by verapamil and CCCP, we quantified fluctuations of the reporter bead during periods of motor arrest. For each motor, we calculated the standard deviation of the bead position in the $x$ and $y$ directions using a 1 s sliding

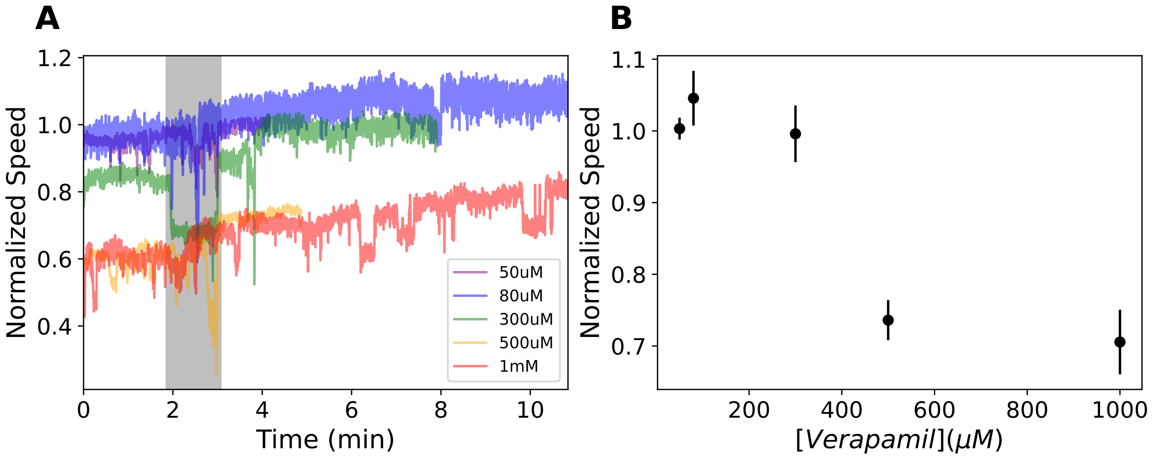


**Figure S2:** BFM speed recovery following verapamil washout. (A) Normalized mean BFM speed before and after verapamil washout for five verapamil concentrations: 50 µM (purple), 80 µM (blue), 300 µM (green), 500 µM (orange), and 1 mM (red) with n = 22, 13, 9, 24, and 33 motors for each concentration, respectively. Grey shaded region indicates the time during which the pump was active; measurements during this interval may be disrupted by flow artifacts. (B) Normalized speed recovery after washout, calculated as the average BFM speed during the last minute following buffer exchange, plotted versus verapamil concentration. Error bars represent standard deviations across individual motors.

window and used $\sqrt{\sigma_x^2 + \sigma_y^2}$ as a measure of the total fluctuation amplitude, normalized to its value during the first second of the recording.

Following CCCP treatment, motors that stopped exhibited substantially larger positional fluctuations than motors stopped by verapamil (Fig. S3). This difference is evident both in individual traces and in the global distributions of fluctuation amplitudes. The increase in fluctuations occurred after motor arrest and persisted throughout the treatment period. Together with the rapid, step-free recovery observed after verapamil washout (Fig. 3A), these observations suggest that verapamil-induced motor arrest places the motor in a mechanical state distinct from that produced by the classical protonophore CCCP.

### 4 Differential Dynamic Microscopy

DDM analysis was performed by using previously described methods [1–3]. We compute the differential intensity correlation function (DICF), defined as the squared modulus of the Fourier transform of the difference of two images separated by a time lag $\tau$, averaged over starting times:

$$\mathrm{DICF}(\vec{q},\tau) = \left\langle \left|\mathcal{F}\left[I(\vec{r},t+\tau) - I(\vec{r},t)\right]\right|^2 \right\rangle_t, \tag{1}$$

where $I(\vec{r},t)$ is the image intensity at position $\vec{r}$ and time $t$, and $\mathcal{F}$ denotes the Fourier transform. Assuming that image intensity is proportional to cell density, the DICF is related to the intermediate scattering function (ISF) through

$$\mathrm{DICF}(q,\tau) = A(q)\left[1 - \mathrm{ISF}(q,\tau)\right] + B(q), \tag{2}$$

where $A(q)$ captures the sample- and optics-dependent amplitude and $B(q)$ accounts for camera noise. Under the assumption of isotropy, both DICF and ISF depend only on $\tau$ and on the magnitude $q = |\vec{q}|$.

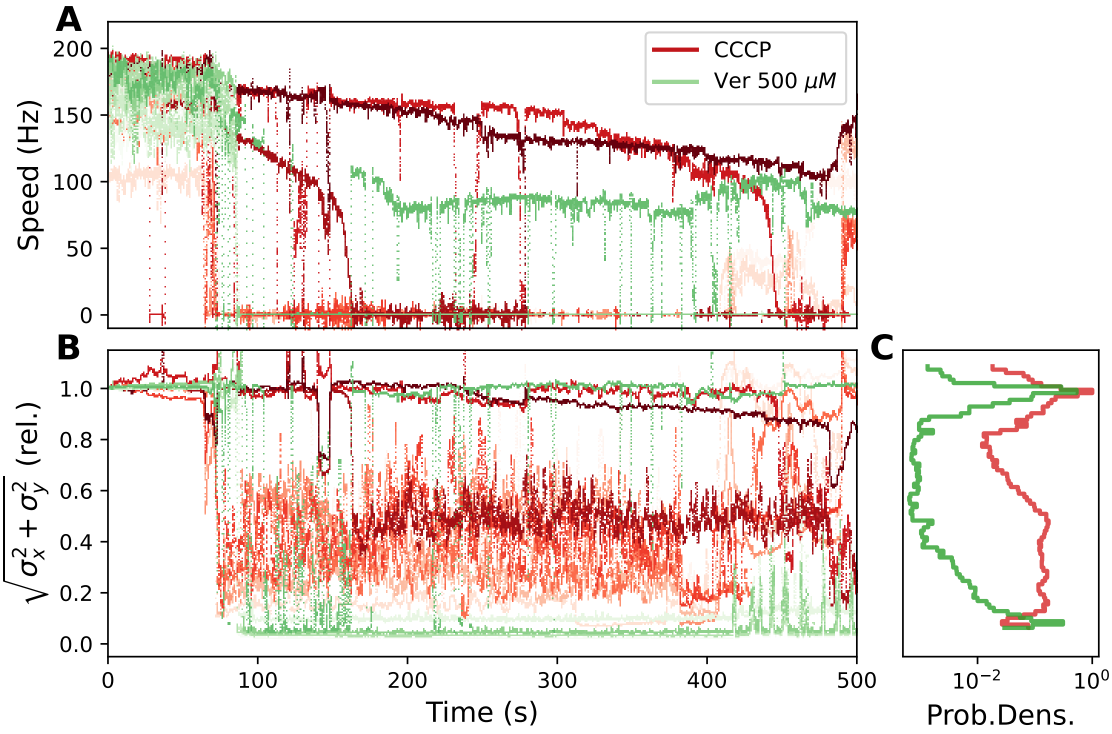


**Figure S3:** Bead fluctuations differ between motors arrested by CCCP and verapamil. (A) Rotational speed traces of individual motors treated with either 20 µM CCCP (red) or 500 µM verapamil (green). Treatment begins at approximately $t = 90$ s. Several motors stop during the recording. (B) Time evolution of the normalized bead fluctuation amplitude, $\sqrt{\sigma_x^2 + \sigma_y^2}$, where $\sigma_x$ and $\sigma_y$ are the standard deviations of the bead position calculated over a 1 s sliding window. Fluctuation amplitudes were normalized to their values during the first second of the recording. Motors arrested by CCCP exhibit substantially larger fluctuations than motors arrested by verapamil. (C) Log-scaled probability density distributions of the normalized fluctuation amplitude shown in (B).

For smooth-swimming cells, we model the ISF using a mixture of diffusive and swimming contributions:

$$ISF(q,\tau) = (1-\alpha)e^{-q^2 D\tau} + \alpha e^{-q^2 D\tau} \int_v P(v) \frac{\sin(qv\tau)}{qv\tau}\, dv, \tag{3}$$

where $\alpha$ is the fraction of motile cells, $D$ is an effective translational diffusion coefficient [4], and $P(v)$ is the Schulz distribution,

$$P(v) = \frac{v^Z}{Z!} \left(\frac{Z+1}{\bar{v}}\right)^{Z+1} \exp\left[-\frac{v(Z+1)}{\bar{v}}\right], \tag{4}$$

with mean speed $\bar{v}$ and shape parameter $Z$ related to the standard deviation $\sigma = \bar{v}(Z+1)^{-1/2}$.

When swimming speeds and motile fractions are low, the ballistic and diffusive contributions to the ISF decay overlap in timescale, resulting in poorly separated relaxation modes [4]. Under these conditions, directly fitting all parameters $(D, \alpha, \bar{v}, Z)$ at each $q$ leads to substantial parameter degeneracies and non-physical solutions. In particular, unconstrained optimization drives $Z$ to its lower bound, producing unrealistically broad velocity distributions and artificially small swimming speeds. This also affects the inferred values of $D$ and

$\alpha$ (as shown in Fig. S4A, continuous lines), which can be overestimated despite visually acceptable fits (Fig. S4A).

To obtain physically meaningful fits, we adopted a constrained fitting strategy. For each $q$ (in the range 1.1–1.9 $\mu m^{-1}$), we fixed the shape parameter to $Z = 2$, ensuring a reasonable speed-distribution width, and explored a grid of prescribed mean speeds $\bar{v}$. For each trial $\bar{v}$, the remaining parameters $D$ and $\alpha$ were fitted by nonlinear least-squares minimization. Fit quality was assessed from the mean squared residual, normalized by the number of degrees of freedom,

$$\chi^2 = \frac{1}{N-p} \sum_{i=1}^{N} \left[ \frac{I_{\mathrm{exp}}(t_i) - I_{\mathrm{fit}}(t_i)}{\sigma_i} \right]^2, \tag{5}$$

where $N$ is the number of time points, $p$ the number of fitted parameters, and $\sigma_i$ the standard deviation of the residuals. The fitting procedure was repeated over a range of fixed mean velocities $\bar{v}$ from 2 to 25 $\mu m\, s^{-1}$, and the optimal value $v$ was selected as the one minimizing the average $\langle \chi^2 \rangle$ across all $q$. This approach reduces the velocity-distribution artifacts that arise when all parameters are free and yields more physically consistent trends. Consistent with earlier work [2, 4], the fitted diffusion coefficient increases with the motile fraction $\alpha$, reflecting the contribution of advective fluctuations generated by nearby swimmers.

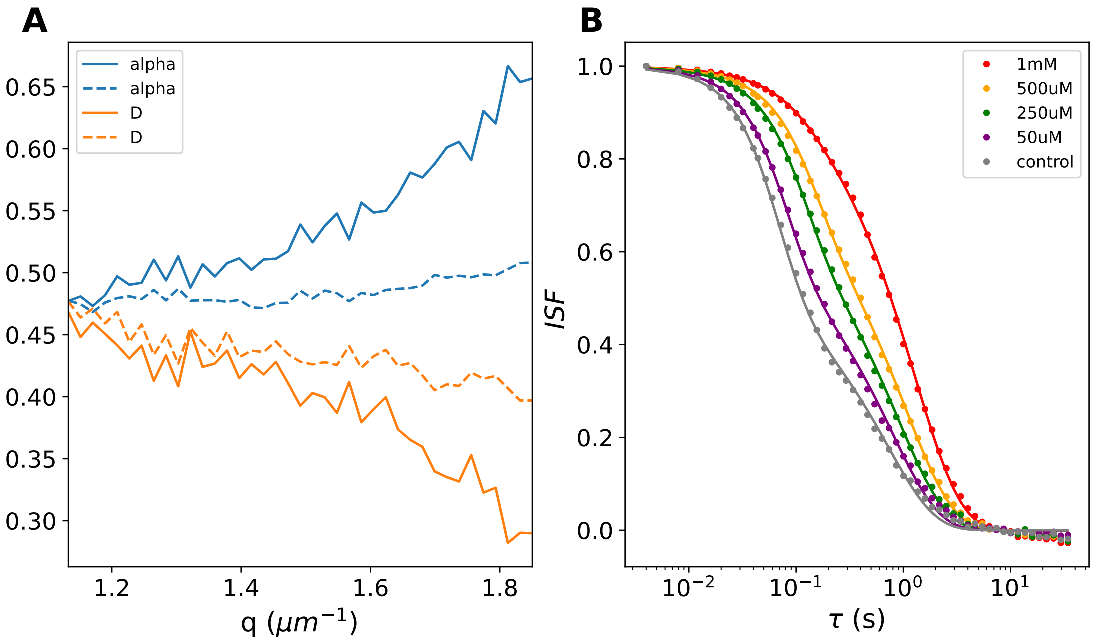


**Figure S4:** Comparison of DDM fitting approaches and ISF behavior under verapamil treatment. (A) Fraction of motile cells ($\alpha$) and effective diffusion coefficient ($D$) obtained using standard DDM fits (solid lines) versus fits in which the Schulz-distribution shape parameter is fixed to $Z = 2$ (dashed lines). Both quantities are shown as a function of the scattering wave vector $q$. Fixing $Z$ suppresses the degeneracy that leads to inflated $\alpha$ and $D$ values in the standard fit. (B) Intermediate scattering functions (ISFs) at fixed wave vector $q$ for cells exposed to different verapamil concentrations. Increasing verapamil concentrations shift the ISF toward slower decay times, indicating a reduction in swimming speed.

#### 5 Antimicrobial susceptibility assay

Minimum inhibitory concentrations (MICs) of verapamil were determined using a standard broth microdilution method. Stationary-phase cultures of *E. coli* were diluted into fresh LB medium and dispensed into 96-well plates containing a two-fold dilution series of verapamil. Each well contained a total volume of 200 µL. $OD_{600}$ was monitored over 15 h in a microplate reader at 35 °C with shaking. The MIC was defined as the lowest concentration at which no net growth was detected relative to the untreated control.

For MT03, growth inhibition was first observed at 10 mM verapamil (Fig. S5A). In contrast, the Δ*tolC* mutant exhibited complete growth inhibition at concentrations as low as 625 µM (Fig. S5B), consistent with an important role for TolC in verapamil tolerance.

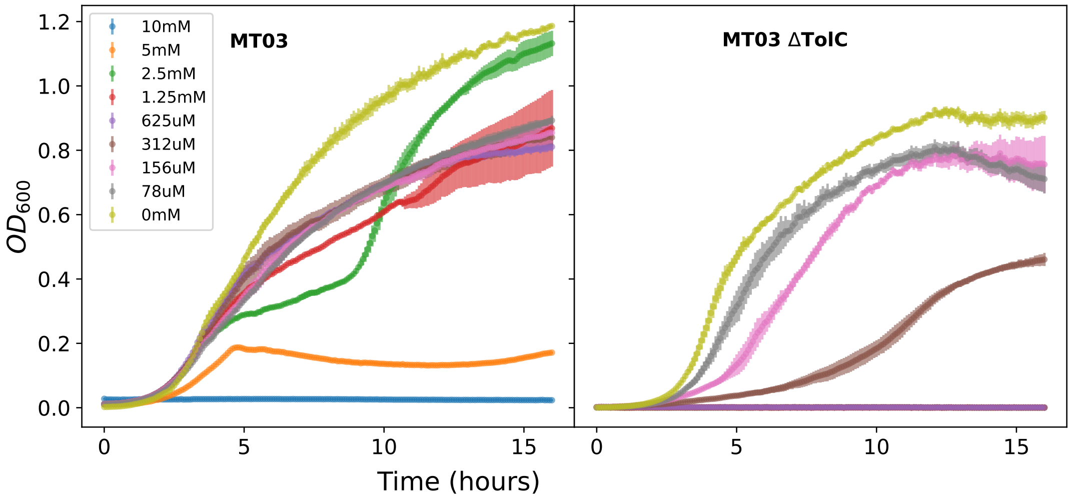


**Figure S5:** Antimicrobial susceptibility assay for MT03 and MT03 Δ*tolC*. Optical density at 600 nm ($OD_{600}$) was monitored over 15 h for cells grown in LB containing increasing concentrations of verapamil. Growth of MT03 was inhibited at 10 mM verapamil, whereas the MT03 Δ*tolC* strain showed complete growth inhibition at concentrations of 625 µM and above. $OD_{600}$ was recorded every 5 min. Shaded regions indicate the standard deviation across three technical replicates.

To assess culturability, ten-fold serial dilutions of the samples were prepared in MB and spotted onto LB agar plates using a 96-pin replica plater (Sigma R2383). Plates were incubated overnight at 30 °C. For MT03, we examined samples grown in 0 mM, 5 mM, 10 mM, and 20 mM verapamil, along with a no-cell control. For MT03 Δ*tolC*, we plated samples from 0 mM, 625 µM, and 1.25 mM verapamil, plus a no-cell control. These concentrations correspond to regimes in which growth was strongly inhibited in bulk assays.

For MT03, approximately $2.5 \times 10^{11}$ CFU/mL were recovered in the absence of verapamil, and $\sim 5 \times 10^6$ CFU/mL at 10 mM, indicating that cells remain culturable under conditions where growth is inhibited. No colonies were observed at 20 mM. In contrast, for the MT03 Δ*tolC* strain, no colonies were recovered at either 625 µM or 1.25 mM verapamil, consistent with its increased sensitivity. These results suggest that verapamil is primarily bacteriostatic at intermediate concentrations in wild-type cells but becomes bactericidal at higher concentrations, whereas loss of TolC shifts this transition to substantially lower concentrations.

### 6 Internal pH measurements

#### Calibration

Ratiometric pHluorin calibration was performed following established protocols [5–7]. Calibration measurements were carried out in parallel with verapamil experiments using dedicated wells. Cells were first incubated in motility buffer (MB) for 30 min to allow adaptation to the low-nutrient conditions, then transferred into buffers containing either 20 µM CCCP or 40 mM PBMH (potassium benzoate and methylamine hydrochloride), which are commonly used to collapse the transmembrane pH gradient. External pH values were adjusted over the range 5.5–9.0. All solutions, including verapamil and calibration buffers, were dispensed simultaneously across the microplate using a 96-channel pipetting robot (Integra Mini 96).

Fig. S6A shows the time evolution of the fluorescence ratio ($I_{395}/I_{475}$; emission at 515 nm) for CCCP (top) and PBMH (bottom). The first three time points (corresponding to the initial 15 min) were used to construct calibration curves of fluorescence ratio versus external pH (Fig. S6B). The data were fitted with the standard sigmoidal function:

$$y = \frac{a_1 e^{k(pH - pH_0)} + a_2}{e^{k(pH - pH_0)} + 1} \tag{6}$$

where $a_1$, $a_2$, $k$ and $pH_0$ are free parameters [6]. Consistent with previous work showing that PBMH most accurately reproduces the *in vitro* calibration of purified pHluorin [6], we used the PBMH-derived curve for all intracellular pH estimates reported in the main text.

Interestingly, we observed that the fluorescence ratio obtained with CCCP at alkaline pH was systematically lower than expected based on prior studies [6]. When interpreted using the PBMH calibration, these measurements suggest that CCCP leads to cytoplasmic acidification rather than a simple collapse of $\Delta pH$, consistent with the mechanism proposed by Plavšek *et al.* [8].

#### Intensity ratio dependence on the incubation time

Fig. S7A shows the evolution of the fluorescence ratio in cells treated with 250 µM verapamil over time for different external pH conditions. A gradual, pH-dependent drift in the fluorescence ratio is observed, leading to systematic differences between early and late time points. These trends complicate quantitative interpretation at longer times. The origin of this drift is unclear, but may reflect pH-dependent changes in verapamil partitioning or cellular adaptation. The intracellular pH values reported in the main text are calculated from measurements within the initial 15 min.

### 7 LIVE/DEAD fluorescence for viability estimation

To calibrate the green/red fluorescence ratio as a quantitative proxy for bacterial viability, we performed LIVE/DEAD BacLight assays on mixtures of live and dead *E. coli*. Cell death was induced by either heat shock or isopropanol treatment to generate suspensions with defined fractions of non-viable cells. For each mixture, fluorescence was measured at 530 nm (SYTO9; green) and 630 nm (propidium iodide; red), and the green/red ratio was computed. Calibration curves showed an approximately linear relationship between this ratio and the known proportion of live cells (Fig. S8). Viability estimates reported in the

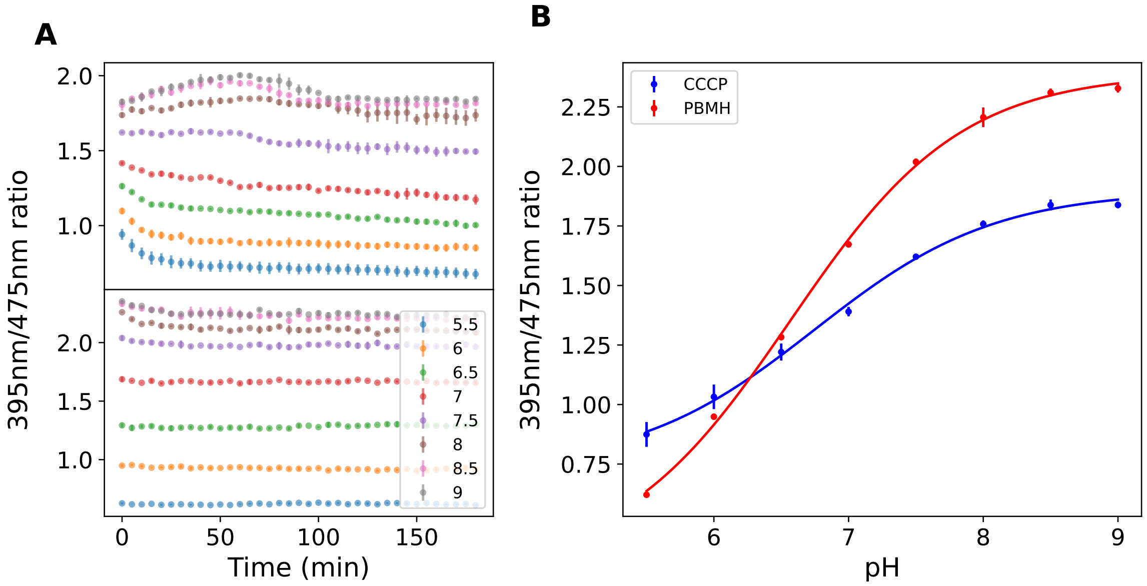


**Figure S6:** pHluorin calibration using CCCP and PBMH. (A) Time course of the pHluorin fluorescence ratio (395 nm / 475 nm excitation; 515 nm emission) for cells incubated at external pH values from 5.5 to 9.0. Two Δ pH–collapsing agents were used: 20 µM CCCP (top) and 40 mM PBMH (bottom). (B) Fluorescence ratio as a function of external pH, computed from the first three time points in (A) (corresponding to the first 15 min). The PBMH data were fitted with a sigmoidal function and used as the calibration curve for determining intracellular pH in the main text. Points represent the mean of three biological replicates; error bars show the standard deviation.

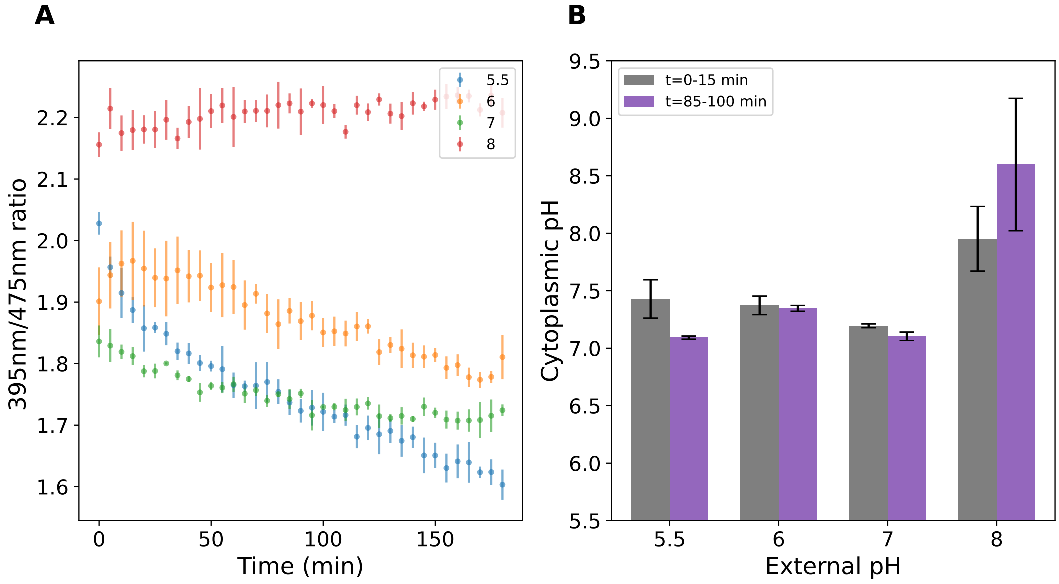


**Figure S7:** Time dependence of pHluorin signals in the presence of 250 µM verapamil. Cells were incubated in buffers with external pH values from 5.5 to 8.0. (A) Time course of the pHluorin fluorescence ratio (395 nm/475 nm excitation; 515 nm emission) at each external pH. (B) Cytoplasmic pH estimated from the calibration curve using the average fluorescence ratio over the first 15 min (gray) and over the 85–100 min interval (purple).

main text were obtained using the calibration derived from heat shock-induced lethality (Fig. S8D).

Fig. S9A–B shows the green and red fluorescence intensities used to compute the ratios in Fig. S8D. Figs. S9D–E show fluorescence intensities for cells treated with 10 µM or 20 µM CCCP. Under these conditions, the green/red fluorescence ratio increases relative to untreated cells (Fig. S9D–F), suggesting an apparent increase in viability. This behavior is counterintuitive, as CCCP collapses the proton motive force without directly compromising membrane integrity, and would therefore not be expected to increase the green/red fluorescence ratio. One possible explanation is that the LIVE/DEAD BacLight assay is sensitive not only to membrane integrity but also to membrane energetics. For example, Kirchhoff *et al.* showed that membrane potential can influence PI uptake, leading to increased red fluorescence even in viable cells [9]. Conversely, collapse of membrane potential by CCCP may reduce PI uptake, resulting in a lower red signal even when membranes are compromised. CCCP may also inhibit efflux activity, allowing SYTO9 to accumulate and further increase the green signal. Together, these effects could bias the green/red fluorescence ratio upward, producing an apparent increase in viability despite energetic disruption, and highlight the need for caution when interpreting these dyes under conditions where membrane potential is perturbed.

### 8 Steady-state motor speeds in Δ*tolC* and wild-type strains

We found that the steady-state rotational speeds of the MT03 Δ*tolC* strain were comparable to those of MT03 (Fig. S10). This contrasts with prior reports of enhanced motility in Δ*tolC* mutants based on soft-agar assays, which quantify radial spreading rather than single-cell swimming speed [10], as well as analogous observations in *Sinorhizobium meliloti* [11]. Related regulatory effects have been reported for other components of the AcrAB–TolC efflux system. For example, deletion of *acrB* upregulates expression of flagellar and motility genes and enhances spreading in soft agar [12], while deletion of *acrR*, a repressor of *acrAB*, produces similar phenotypes through derepression of the *flhDC* master regulator [13, 14].

However, increased spreading on soft-agar plates does not necessarily reflect an increase in swimming speed. Instead, it can arise from altered chemotactic behavior, such as longer run durations, which expand the apparent motility zone even in the absence of faster swimming. This interpretation is consistent with recent work showing that, although regulatory perturbations can increase flagellar number, hydrodynamic constraints limit the extent to which per-cell swimming speed can increase [15, 16]. Our direct measurements of flagellar motor rotation therefore indicate that the previously reported motility differences in Δ*tolC* cells likely reflect changes in chemotaxis or population-scale spreading dynamics rather than intrinsic changes in swimming speed.

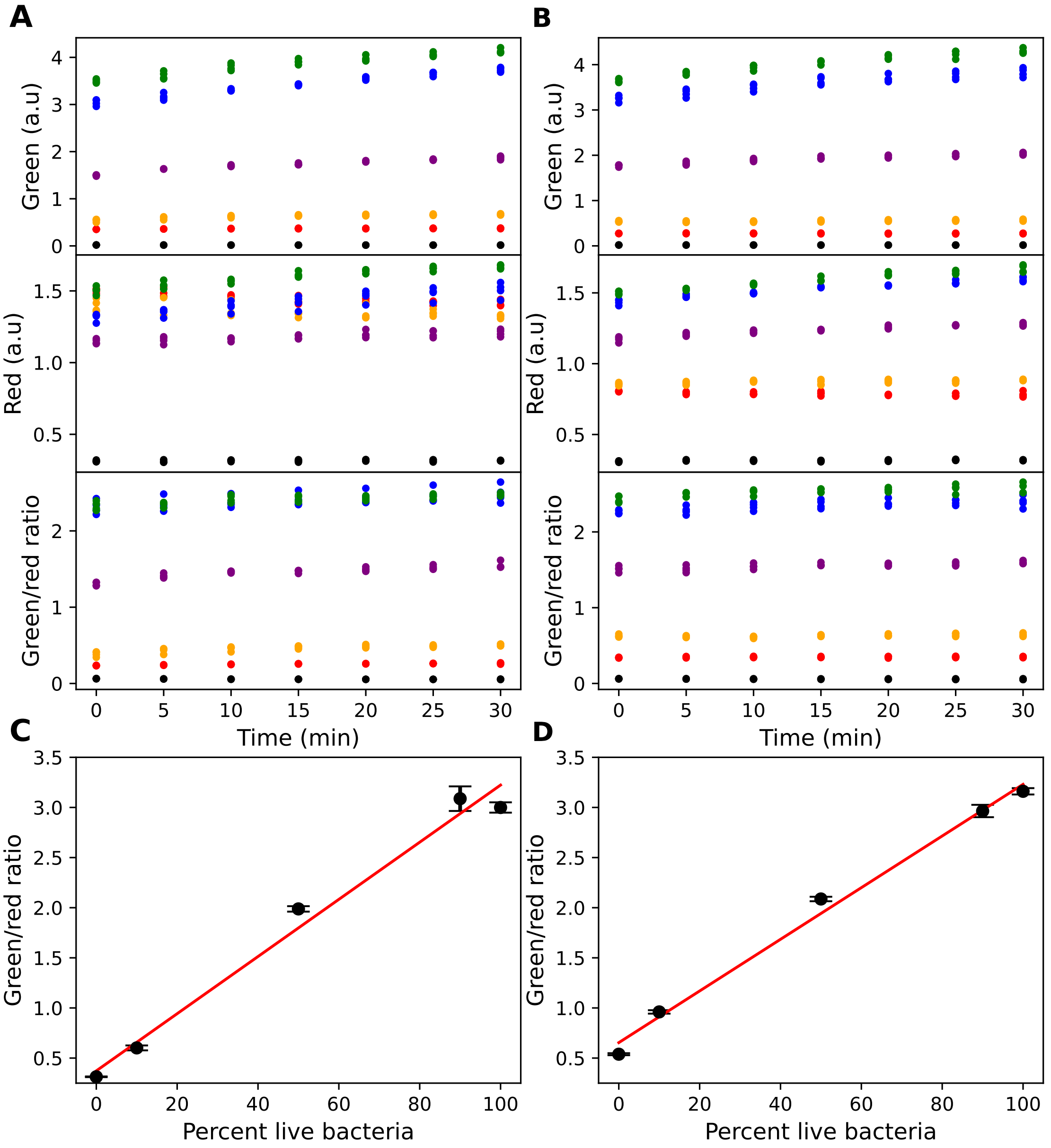


**Figure S8:** Calibration of BacLight fluorescence ratio with live/dead mixtures. (A–B) Green fluorescence (top), red fluorescence (middle), and green/red fluorescence ratio (bottom) for mixtures of live and killed *E. coli* using isopropanol treatment (A) or heat shock (B). Live-cell fractions are: 0% (red), 10% (orange), 50% (purple), 90% (blue), and 100% (green). Black points correspond to dye-only controls. (C–D) Green/red fluorescence ratio at 15 min as a function of the percentage of live cells for isopropanol-treated (C) and heat-killed mixtures (D). Ratios were background-corrected using no-cell controls, $R = (g - g_0)/(r - r_0)$. Red lines indicate least-squares fits, with $R^2 = 0.99$ for (C) and $R^2 = 0.98$ for (D).

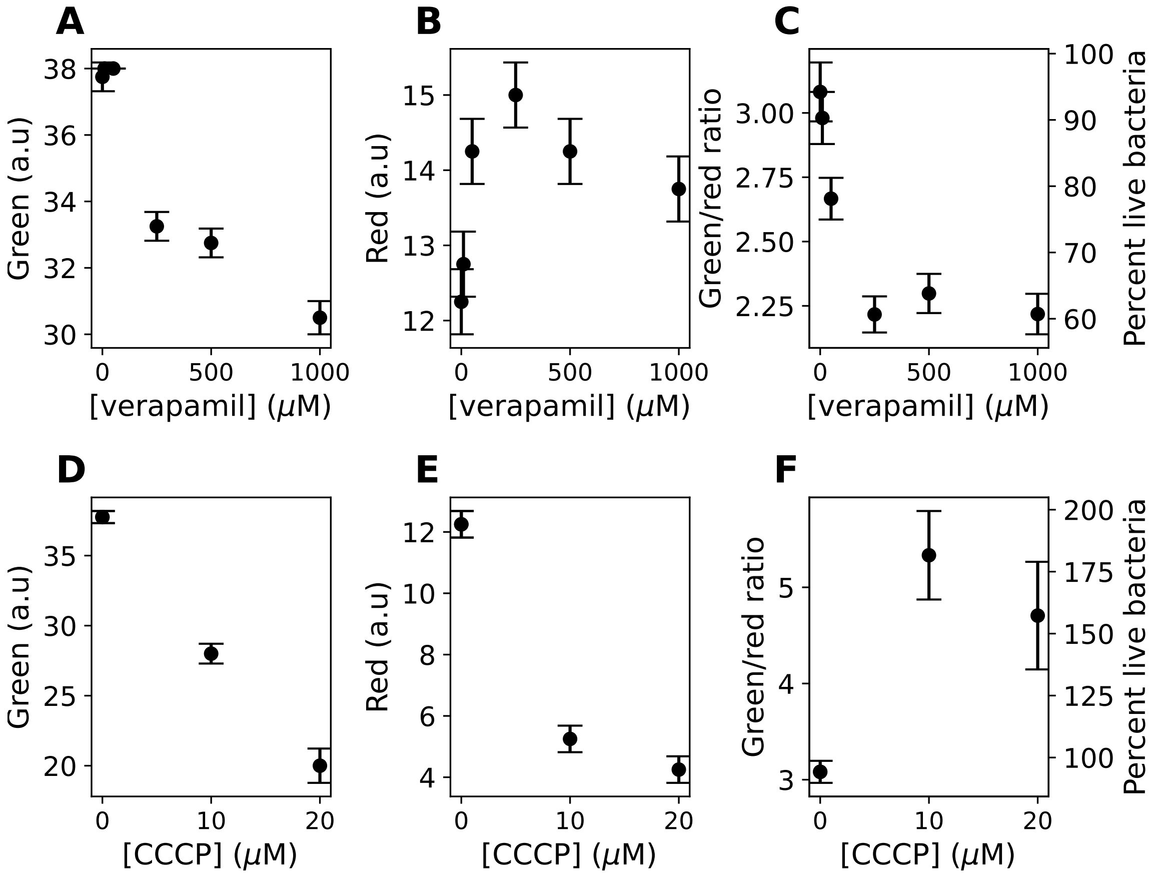


**Figure S9:** Effect of verapamil and CCCP on BacLight fluorescence signals and inferred viability. (A–C) Green fluorescence intensity, red fluorescence intensity, and green/red fluorescence ratio of bacterial suspensions treated with increasing concentrations of verapamil. (D–F) Same as (A–C), but for cells treated with increasing concentrations of CCCP. In (C) and (F), the right y-axis shows the estimated percentage of live cells, calculated from the linear calibration between green/red fluorescence ratio and the fraction of live cells (see Fig. S8). Each point represents the mean of three independent measurements; error bars represent the standard deviation of the mean (A–B, D–E) or are derived from error propagation analysis (C, F).

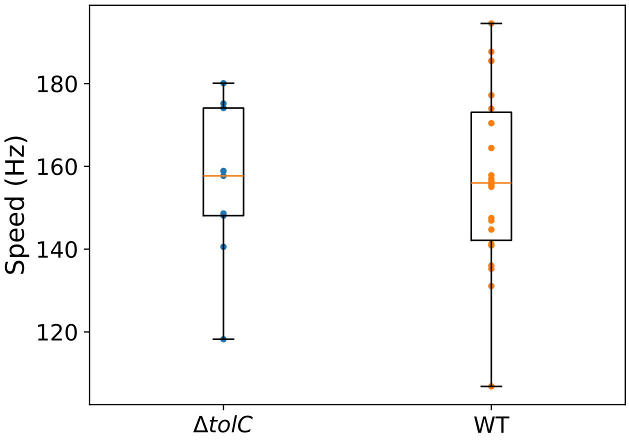


**Figure S10:** Steady-state motor speeds of the $\Delta tolC$ and wild-type strains. Boxes indicate the interquartile range, with the median shown as a horizontal line. Whiskers extend to the most extreme data points within 1.5 times the interquartile range. Sample sizes: $\Delta tolC$, $n = 8$ (blue); wild type, $n = 22$ (orange).
